# Bioinformatic analysis of B and T cell epitopes from SARS-CoV-2 Spike, Membrane and Nucleocapsid proteins as a strategy to assess possible cross-reactivity between emerging variants, including Omicron, and other human coronaviruses

**DOI:** 10.1101/2022.02.16.480759

**Authors:** Diana Laura Pacheco-Olvera, Stephanie Saint Remy-Hernández, María Guadalupe García-Valeriano, Tania Rivera-Hernández, Constantino López-Macías

## Abstract

The COVID-19 pandemic caused by SARS-CoV-2 produced a global health emergency since December 2019, that up to the end of January 2022 had caused the death of more than 5.6 million people worldwide. Despite emergence of new variants of concern, vaccination remains one of the most important tools to control the pandemic. All approved vaccines and most of the vaccine candidates use the spike protein of the virus as a target antigen to induce protective immune responses. Several variants of the virus present key mutations in this protein which render the virus, at different rates, to evade the neutralizing antibody response. Although experimental evidence suggests that cross-reactive responses between coronaviruses are present in the population, it is unknown which potential antigens shared between different coronaviruses could be responsible for these responses. This study provides predictions of new potential B and T cell epitopes within SARS-CoV-2 Spike (S), Membrane (M) and Nucleocapsid (N) proteins together with a review of the reported B epitopes of these proteins. We also analyse amino acid changes present in the epitopes of variants of concern (VOC) and variants being monitored (VBM), and how these might affect the immune response, as these changes may alter the peptides’ immunogenicity index and the antigen presentation by related HLA alleles. Finally, given these observations, we performed an identity analysis between the repertoire of potential epitopes of SARS-CoV-2 and other human coronaviruses to identify which are conserved among them.

The results shown here together with the published experimental evidence, allow us to support the hypothesis that antibody and T cell cross-reactive responses to common coronaviruses epitopes, could contribute to broaden the protective response to SARS-CoV-2 and its variants. This evidence could help not only to understand cross-reactive responses among coronaviruses but also contribute to elucidate their role in immunity to SARS-CoV-2 induced by infection and/or vaccination. Finally, these findings could promote targeted analysis of antigen-specific immune responses and might orient and drive the rational development of new SARS-CoV-2 vaccines including candidates that ideally provide “universal” protection against other coronaviruses relevant to human health.

## Introduction

Over the past 30 years, three coronaviruses, SARS-CoV, MERS-CoV, and SARS-CoV-2, have evolved and adapted to cause severe respiratory diseases and spread among the human population. On the other hand, there are four seasonal coronaviruses, HKU1, NL63, 229E, and OC43, which cause mild respiratory diseases in humans^1^. Today, we are still in the battle to stop the pandemic caused by SARS-CoV-2, responsible for 313 million confirmed cases as of January 2022 and more than 5.6 million deaths worldwide^2,3^. Unfortunately, despite the effectiveness of vaccines, cases continue to increase due to constant changes occurring in the genome of SARS-CoV-2 that have affected mainly the gene encoding for the spike protein (S) used in the current approved vaccines. Mutations in this antigen which have been identified in specific regions of S protein are important for the interaction with angiotensin-converting enzyme 2 (ACE2), the host cell-specific receptor, render some variants with changes to improve infectivity, transmissibility, and resistance to neutralizing antibodies induced by vaccines^4^.

Currently, the World Health Organization (WHO) recognizes five SARS-CoV-2 variants of concern (Alpha, Beta, Gama, Delta, and Omicron) and seven variants of interest (Epsilon, Eta, Iota, Kappa, Lambda, Theta, and Zeta)^5^ and this number is likely to increase in the future as long as the virus keeps actively circulating. Variant B.1.1.7 (Alpha) was identified in the United Kingdom (UK) in September 2020, this variant is highly transmissible thanks to several mutations in the S protein that help the virus adhere more strongly to human cells and can help infected cells create new S proteins more efficiently. In addition, H69-V70 and Y144/145 deletions alter the form of the S protein which may help the virus to evade certain antibodies^6^. The level of protection against Alpha variant conferred by vaccination, is similar to the observed in clinical trials, showing a modest decrease in neutralizing activity of samples from vaccinated people, while maintaining the same level of protection against severe disease^7,8^. Variant B.1.351 (Beta), first detected in South Africa, showed approximately 20-fold increase in affinity for ACE2 compared to the receptor-binding domain (RBD) of the Wuhan variant, making it more transmissible^9^ and able to escape the neutralization of monoclonal antibodies due to the E484K mutation^10^.

Laboratory data showed a moderately reduced efficacy against symptomatic disease with this variant, but high levels of efficacy (97%) against severe disease in people vaccinated with the BNT162b2 vaccine (Pfizer/BioNTech) and 51% efficacy for the NVX-CoV2373 (Novavax) vaccine against this variant^11,12^. Another extremely infectious variant is P.1 (Gamma), firstly reported in Brazil in mid-2020. This variant has severely increased the number of infections in this country collapsing the Brazilian health system^13^. However, high neutralization titres against this variant have been observed in sera from people vaccinated with BNT162b2^14^. Variant B.1.617.2 (Delta) was detected in India in December 2020, mutations in this variant cause increased replication, leading to high viral loads and increased transmission. Two doses of the BNT162b2 vaccine or ChAdOx1 nCoV-19 (AstraZeneca) vaccine within the previous 39 days are effective to prevent (88% and 67%, respectively) symptomatic disease induced by delta variant ^15^. Similarly, lineages B.1.427 and B.1.429 of the Epsilon variant detected in California, United States, have significant mutations in S protein (S13I, W152C, and D614G) and RBD (L452R). Another variant, B.1.525 (Eta), identified in the UK and Nigeria, presents several significant mutations in S protein (A67V, 69del, 70del, 144del, D614G, Q677H, and F888L) and RBD (E484K). Some of these mutations are associated with increased infectivity, transmission, and a reduction in neutralization. Variant B.1.617.1 (Kappa) circulating in India, has important mutations in the S protein (T95I, D614G, E154K, P681R, G142D, and Q1071H) and particularly crucial in RBD (E484K and L452R). In the case of the variants P.2 (Zeta) and P.3 (Theta) identified in Brazil and Japan respectively, they present some similar and important mutations in S protein (D614G and V1176F) and RBD (E484K), which influence the reduction of neutralization by antibodies and an increase in its ability to cause reinfection^16–19^. Finally, in November 2021 the Omicron variant (B.1.1.529), which was first detected in South Africa, harbours 30 mutations in the S protein, amongst them an insertion of three amino acids (EPE 214) and other changes such as V143/Y145^20^. Some of these mutations have been previously reported in the Alpha, Beta, Gamma, and Delta variants, as well as Kappa, Zeta, Lambda, and Mu variants, and have been associated with increased transmissibility and evasion of immune responses^21^. Omicron has been the variant in which the greatest number of mutations have been described so far; within the M protein three mutations: D3G, Q19E and A63T, have been reported, whereas the N protein presents three substitutions and one deletion of three residues^20^.

Protective immunity against SARS-CoV2 involves both antibody and T cell immune responses; adaptive immune responses mediated by CD8+ and CD4+ T cells are essential to promote an efficient B cell and antibody responses as well for the elimination of cells infected by the SARS-CoV-2 virus, whereas B cell and antibody responses are important to generate neutralizing antibodies to control the viral infection. Antibody and T cell responses are directed to several SARS-CoV-2 components such as M, N and S protein among others^22^. S protein has been utilised as the main target to induce protective antibody and T cell responses, nevertheless, the contribution of the immune responses elicited by other viral antigens remains poorly explored^23^. SARS-CoV-2 epitopes targeted by CD4+ and CD8+ T cells are different between disease states (mild to severe cases), where mild cases respond to a wider repertoire of epitopes among the entire viral proteome, severe cases respond to a reduced repertoire ^23,24^. On the other hand, there is evidence of a cross-reactive response with other human coronaviruses, potentially explained by homology between the sequences of these coronaviruses and SARS-CoV-2. These T-cell cross-reactive responses to SARS-CoV-2^25^ could help to explain the different clinical pictures in COVID-19 patients. For antibody responses, it has been observed that during mild disease a diverse neutralizing antibody repertoire is formed, whereas in patients with severe disease T and B cell compartments are compromised and antibody responses showed less diversity^26,27^.

Despite experimental evidence suggests that cross-reactive responses between coronaviruses are present amongst the population, little information is available about particular epitopes that might drive these responses. Using web-based bioinformatic tools we have been able to build a repertoire of potential B and T cells epitopes that are found in SARS-CoV-2 and identified those epitopes that are conserved with other coronaviruses relevant to human health. In addition, we present an analysis of mutations found in the S protein of other SARS-CoV-2 variants and show that, while several potential epitopes are affected, the changes in immunogenicity are not necessarily detrimental and on the other hand, there is an ample repertoire of epitopes that remain conserved. The results of this work and together with the experimental evidence that emerges day by day, support the hypothesis that cross-reactive B and T cell responses to coronaviruses common epitopes, could play a key role in the immunity to SARS-CoV2 and its variants induced by infection or vaccination.

## Materials and methods

### Epitope prediction for T and B cells

Genome sequences of human coronaviruses HCoV-HKU1 (HKU), HCoV-NL63 (NL63), HCoV-OC43 (OC43), HCoV-229E (229E), MERS-CoV, SARS-CoV, and SARS-CoV-2 were accessed through NCBI, Genbank access numbers YP_173238.1, YP_003767.1, YP_009555241.1, ABB90529.1, AKN11075.1, AAU04646.1 and QHR63290.2 respectively, multiple S protein sequences were aligned using Clustal Omega^28^

The prediction of linear B cell epitopes was performed through the Immune Epitope Database (IEDB) website (https://www.iedb.org/) using the Bepipred V1.0 and V2.0 linear epitope prediction algorithms^29^. Threshold values of 0.35 were used (corresponding specificity> 0.49 and sensitivity <0.75) and 0.55 (corresponding specificity> 0.817 and sensitivity <0.292) with versions 1.0 and 2.0, respectively. Epitopes were chosen based on their IC50 binding values of <50 (high affinity) and <500 (mean affinity). To determine whether the predicted B cell epitopes were exposed on the surface of the protein, surface accessibility and secondary structure NetSurfP-2.0^30^ was used. For the prediction of the MHC-I restricted T cell epitopes, MHC-I binding predictions (NetMHCpan EL 4.0 method)^31^ and Class I immunogenicity tools^32^ from the IEDB website were used. The selection of T cell epitopes was made based on IC50 affinity scores (<500) and immunogenicity scores (> -1 and <1). For the prediction of the MHC-II restricted T cell epitopes, we used the algorithm 2.22 recommended by IEDB and TepiTool^33^ was used for prediction. Prediction was made for 3 HLA-DR alleles (HLA-DRB1 * 01: 01, HLA-DRB1 * 04: 01, HLA-DRB1 * 07: 01), 8 HLA-DP alleles (HLA-DPA1 * 01: 03 / DPB1 * 03: 01, HLA-DPA1 * 02: 01 / DPB1 * 02: 01,HLA-DPA1 * 02: 02 / DPB1 * 02: 02, HLA-DPA1 * 03: 01 / DPB1 * 23: 01) and 6 HLA-DQ alleles (HLA-DQA1 * 05: 01 / DQB1 * 03: 01, HLA-DQA1 * 01: 01 / DQB1 * 05: 01,HLA-DQA1 * 03: 01 / DQB1 * 03: 02) with a peptide length limit of 15 amino acids and an average consensus percentile of the 20 prediction threshold. The selection of the T cell epitope was made based on IC50 affinity scores (<500)^29,31–33^.

The alleles were used for the prediction given that they are present with a high frequency in the Mexican population and also have a significant frequency worldwide^34–37^. Additional alleles were obtained from the allele frequency database (http://www.allelefrequencies.net/default.asp)^38^, using North American region as the search criteria, and ethnic Mexican groups present in all the State capitals of the Country.

### Analysis of SARS-CoV-2 mutations and clinically significant variants

Sequences of the SARS-CoV-2 genome isolated from different countries were accessed through the GISAID database (https://www.gisaid.org/)^39,40^ filtered from December 2019 to December 30, 2021, using 33 complete SARS-CoV-2 genomes from GISAID data base from different patients around the world (i.e., Australia, Germany, Italy, Mexico, Saudi Arabia, Taiwan, Vietnam, China, Belgium, Ecuador, Malaysia, New Zealand, Singapore, USA, Brazil, Estonia, South Africa, England, India, Botswana, and Hong Kong) with at least one sequence belonging to one of the 10 clades established by GISAID, and a representative sequence of each of the variants of VOC sequences that have been reported. Only the complete genomes (28000 to 30000 bps) were included in the analysis, and the reference sequence of NCBI from Wuhan (ID NC_045512.2) (Supplementary Table 4)

The Transeq EMBOSS tools (https://www.ebi.ac.uk/Tools/st/emboss_transeq/)^28,41^ were used for sequence translation in the 6 nucleotide-to-amino acid reading frames, and the 10 proteins analyzed were manually cured (Spike, Membrane protein and Nucleocapsid). Once the nucleotide sequences were translated into protein, Jalview, Clustal Omega and Blast (https://blast.ncbi.nlm.nih.gov/Blast.cgi)^28,42,43^ were used to perform multiple alignments to compare the various sequences to the reference sequence of the SARS-CoV-2 virus (QHR63290.2).

To provide a graphical representation of epitopes, we used the full-length SARS-CoV-2 S protein structural model (ID: 6VSB_1_1)^44,45^. Graphical representations of M (ID: QHD43419) and N (ID: QHD43423) S protein structures were generated through D-I-TASSER/ C-I-TASER (https://zhanggroup.org/)^46,47^

The structures were built in 3D and analyzed using PyMOL software (Schrödinger LLC. Molecular Graphics System (PyMOL) Version 1.80 LLC, New York, NY 2015^48^. The basic online local alignment search tool (https://blast.ncbi.nlm.nih.gov/Blast.cgi)^43^ was used to evaluate the position of the predicted peptides in the sequence of the proteins analysed in this study.

## Results

### Prediction of B cell linear epitopes

We conducted a literature review of studies that reported the mapping potential epitopes of the spike (S) protein, the membrane protein (M), and nucleocapsid (N) of SARS-CoV-2, the search criteria were using the combination of the words: SARS-CoV-2, epitopes of B and T cells, S/spike protein, M/membrane protein and N/nucleocapsid protein in the search engine PubMed https://pubmed.ncbi.nlm.nih.gov/^49^, Google Scholar https://scholar.google.com/^50^, ScienceDirect https://www.sciencedirect.com/^51^, and COVIDep https://covidep.ust.hk/^52,53^ data base. The review covered reports generated between January 2020 to January 2022.

For the specific case of spike protein, we were able to compile a total of 116 potential B cell epitopes, of which 61 were epitopes previously reported *in silico* (ERIS)^52,54–59^ by other research groups (Supplementary Table 1), on the other hand, we compiled 19 experimentally confirmed epitopes (ECE)^57–60^ (Supplementary Table 3) also reported by others. Using BepiPred 1.0 and 2.0, our study predicted 36 epitopes that were selected based on their score and surface accessibility, and which we have denominated epitopes predicted in this study (EPITS). From these 36 EPITS, 29 had already been reported in studies by other research groups, and 7 were novel reported epitopes (NRE) (Supplementary Table 1). Epitopes from our predictions that overlapped with any ERIS, were fused in a single polypeptide and marked in bold characters, and ECEs are shown in red (Supplementary Table 1 and 3). To visualize the position of the compiled epitopes we used an optimized 3D model of SARS-CoV-2 spike protein in its trimeric conformation (Figure 1A). ERIS are represented in magenta, EPITS are represented in pink, and finally, in purple, we show ECE. B cell epitopes reported *in silico* and described in this study are distributed across all domains of the spike protein. The trimeric model shows several predicted epitopes, from this and other studies, that still lack experimental validation, highlighting the importance of further investigating the immunogenicity of these predicted epitopes. It should be noted that experimentally confirmed epitopes are in the regions comprising the S1 polypeptide, which contains the N-terminal domain (NTD), the receptor-binding domain (RBD) and the receptor-binding motive (RBM)^44,61^.

**Figure 1.**
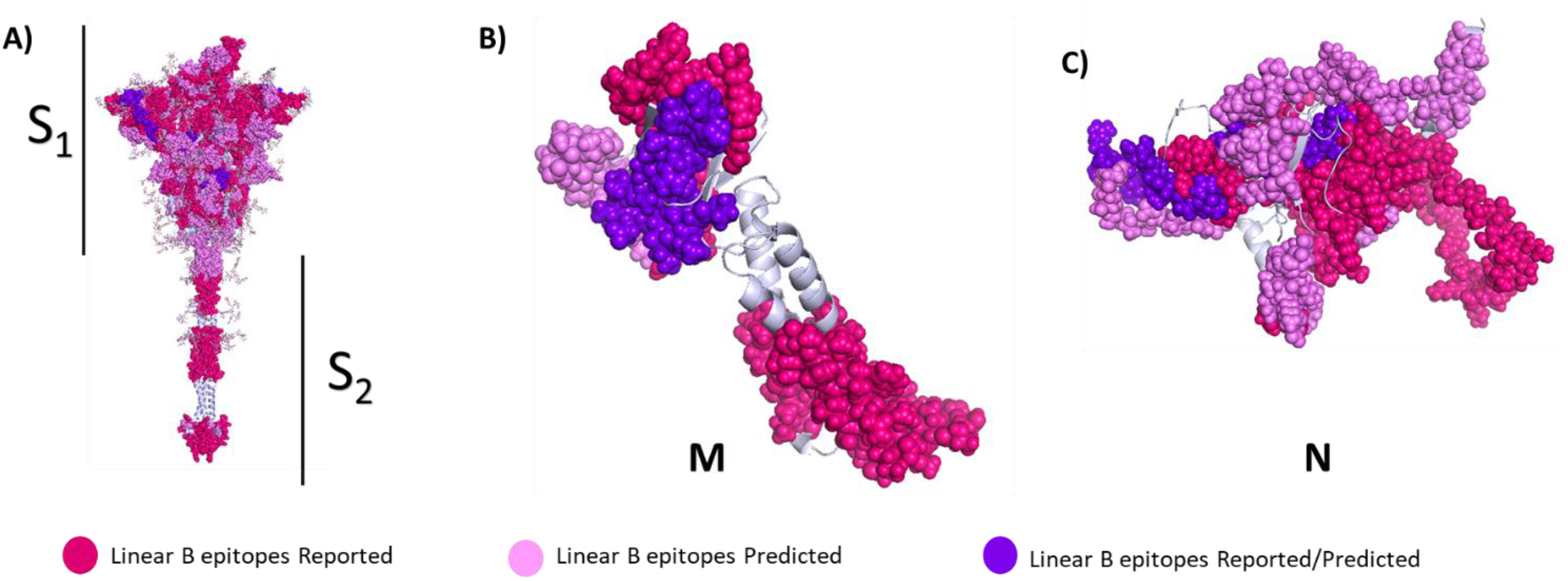
Linear B cell epitopes in spike, membrane and nucleocapsid proteins. 3D graphical representations of (A) spike, (B) membrane and (C) nucleocapsid proteins of SARS-CoV2. Linear B cell epitopes reported by *in silico* studies found in the literature are shown in magenta color, epitopes predicted in this study are shown in pink, and predicted epitopes that have been experimentally confirmed to be immunogenic are shown in purple. X-ray crystallography S protein ID: 6VSB_1_1, M protein ID: QHD43419 and N protein ID: QHD43423.

Following the same workflow that was used for S protein, we performed the analysis for the membrane (M) and nucleocapsid (N) protein sequences. For M protein, we were able to collect a total of 18 potential B cell epitopes (Supplementary Table 1), of which 9 are defined as ERIS^36,52,54,57,60^. In addition, we found in the literature 2 ECE^57^, as shown in Supplementary Tables 1 and 3. Using BepiPred 1.0 and 2.0, we predicted and selected 7 EPITS which were all both, predicted in this study and by other research groups. We did not find any novel epitopes for this protein. For the representation of the compiled epitopes, we use a 3D model of the M protein (Figure 1B), where we highlight the ERIS in magenta, the EPITS in pink, and the ECE are shown in purple. Additionally, we can observe that most of the epitopes reported, predicted and confirmed are located in the C-terminal and the N-terminal domains^62^.

For N protein, we were able to collect a total of 63 potential B cell epitopes; 35 are classified as ERIS^36,52,54,63^, 5 as ECE^63^ (Supplementary Table 1 and 3), and 23 as EPITS, of which 21 were predicted in this study and by other research groups and 2 are NRE (Supplementary Table 1). The epitopes from our predictions that overlapped with ERIS, were fused in a single polypeptide highlighting in bold characters the EPITS and in red the ECE (Supplementary Table 1 and 3). Using the 3D model of N protein (Figure 1C), we were able to locate and visualize the position of each of the ERIS in magenta, EPITS in pink, and the ECE in purple.

### Prediction of T cell epitopes

In total, we identified for S protein 128 CD8 and CD4 T cells epitopes (Figure 2A), 89 for protein M (Figure 2B), and 114 for protein N (Figure 2C). From these, we identified 73 ERIS^36,55,64–69^ for S protein, 77 for M protein^59,60,70–74^ and 98 for N protein^36,52,64,70,72,74,75^ (Supplementary Table 2). In addition, we were able to compile 9 ECE for S protein^64–69,76^, 3 for M protein^64^ and 5 for N protein^64,65,75^. And, we detected 46 EPITS for the S protein, of which 27 are NER: 10 restricted to MHC-I binding, 5 to MHC-II binding, and 12 are predicted to be promiscuous T cell epitopes (presented to both CD4+ and CD8+ T cells). For M protein we identified 9 EPITS of which 2 were NER epitopes, one epitope was restricted for MHC-I and one for MHC-II, while no promiscuous peptides were found (Supplementary Table 2). And finally for N protein we identified 11 EPITS, of which 2 were NER restricted to MCH-I, one NER was classified as promiscuous, and none were MHC-II restricted. For the visualization of epitopes, we used an optimized 3D model of the three proteins (Figure 2), the ERIS are depicted in blue, while EPITS are shown in green and finally CD4+ or CD8+ T cells ECE appear in red. The epitopes described in the literature can be found throughout the structure of spike protein, the predicted epitopes are present throughout the structure except for a notable gap in the transmembrane domain (TD) (Figure 2A). In the case of M protein (Figure 2B) we show that the EPITS are evenly distributed throughout the protein, while ECE are mainly situated in the N-terminal and C-terminal domains. On the other hand, the N protein 3D model shows that the distribution of the epitopes is across the entire protein, while the ECE were mapped mainly in the RNA-binding domain, the binding site and within the dimerization domain (Figure 2C).

**Figure 2.**
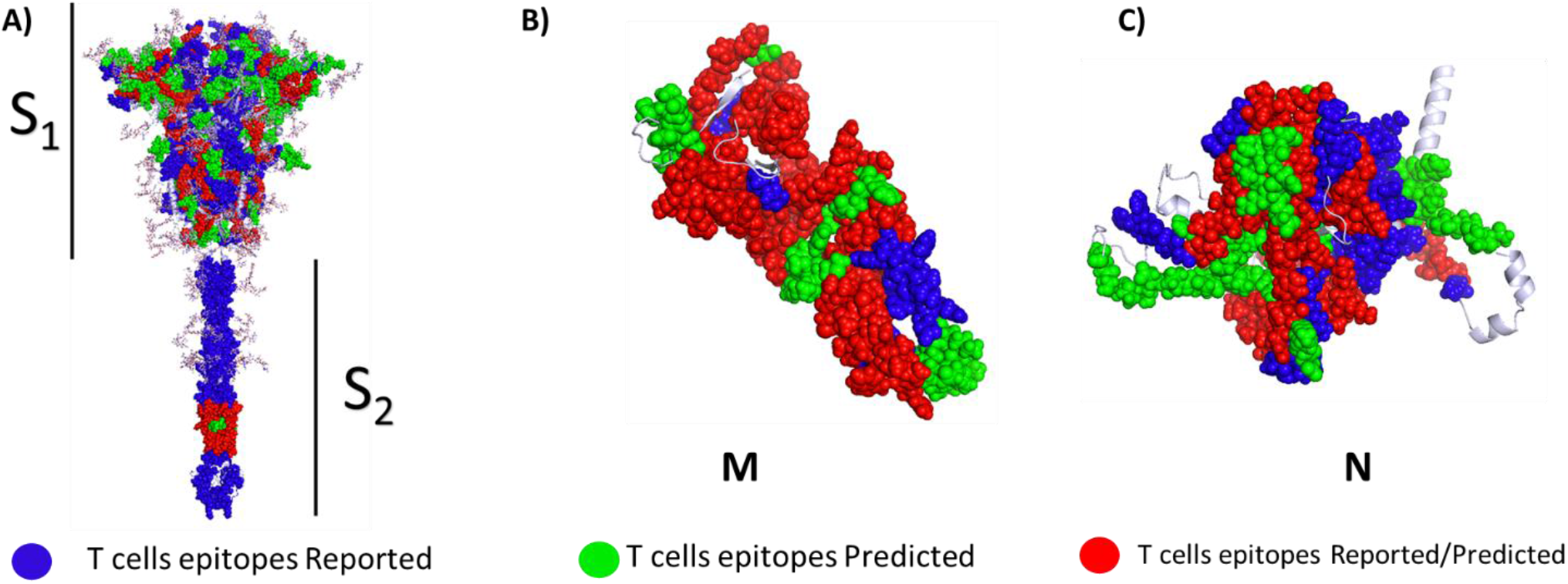
T cell epitopes in spike, membrane and nucleocapsid proteins. 3D graphical representations of (A) spike, (B) membrane and (C) nucleocapsid proteins of SARS-CoV2. T cell epitopes reported by *in silico* studies found in the literature are shown in blue color, epitopes predicted in this study are shown in green, and predicted epitopes that have been experimentally confirmed to be immunogenic are shown in red. X-ray crystallography S protein ID: 6VSB_1_1, M protein ID: QHD43419 and N protein ID: QHD43423.

### Mutations present in SARS-CoV-2 clinically significant variants

To map the mutations reported for the Alpha, Beta, Gamma, Delta and Omicron variants of concern, we used an optimized 3D model of the S protein, where the mutations that have been reported are shown in red. In the model are also depicted the EPITS, CD8+ T cells epitopes are shown in blue, CD4+ T cells epitopes in green, and linear B cell epitopes in purple (Figure 3). This model shows that mutations in variants are mainly located in the S1 polypeptide region, leaving the epitopes that are positioned in other regions of the S protein intact.

**Figure 3.**
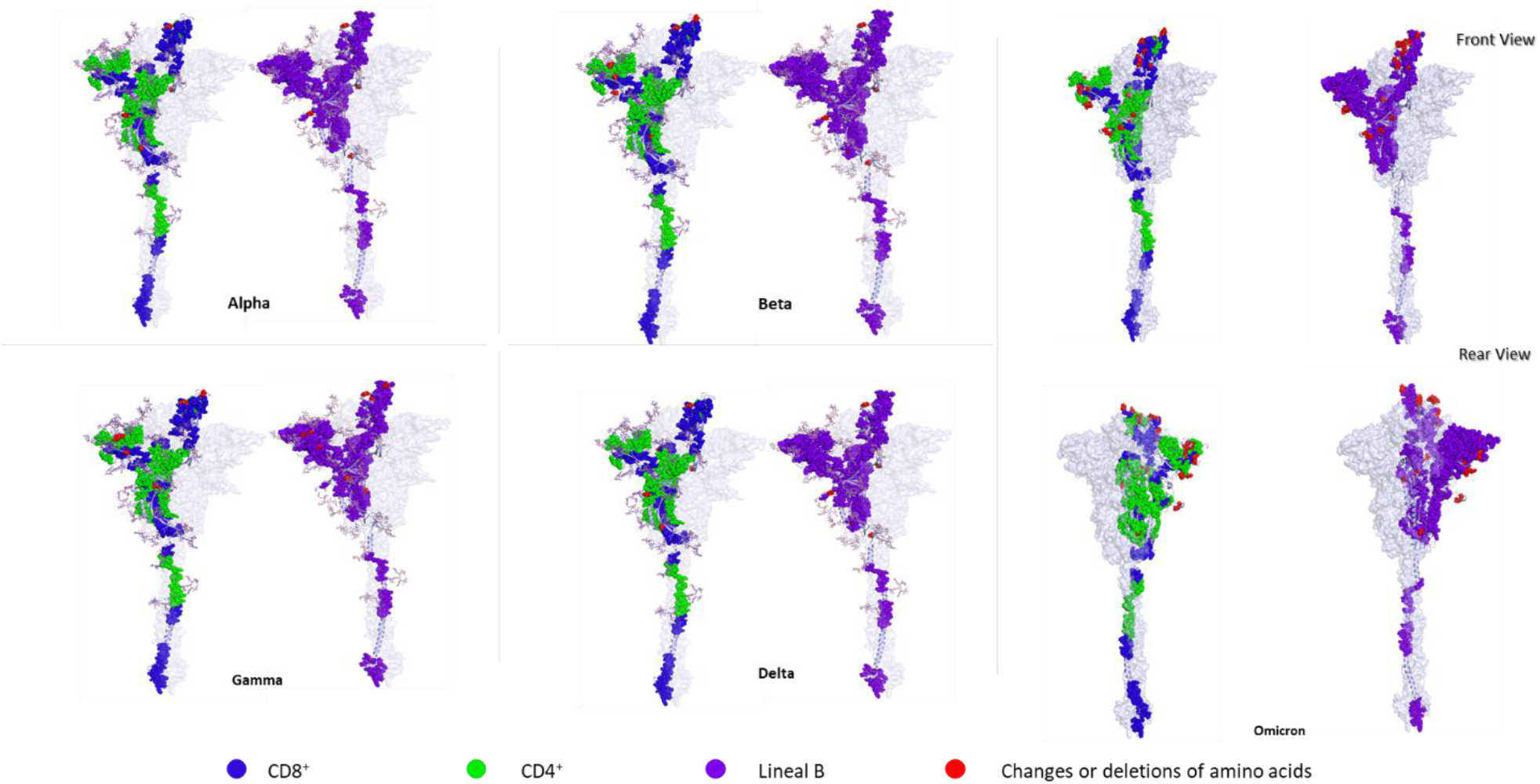
Amino acid changes in SARS-CoV-2 variants being monitored (Alpha, Beta and Gamma) and variants of concern (Delta and Omicron). 3D graphical representations of the spike protein of SARS-CoV2 where changes or deletions in specific amino acids are shown in red for Alpha, Beta, Gamma, Delta and Omicron variants. CD8+ T cells epitopes are shown in blue, CD4+ T cell epitopes are shown in green, and B cell epitopes are shown in purple. All epitopes compiled in this work (ER, EP and ED) are shown. Front and rear views of the Omicron variant are presented to highlight the high number of amino acid changes along the protein sequence. X-ray crystallography S protein ID: 6VSB_1_1.

To evaluate epitope modifications in different SARS-CoV2 isolates from different geographical regions, a multiple sequence alignment analysis of the spike, membrane and the nucleocapsid proteins of SARS-CoV-2 was carried out (Supplementary Tables 3, 5 and 6). This allowed us to identify epitopes that vary between the reference sequence with respect to other selected sequences. We detected modifications in the sequences of the 3 evaluated proteins (S, M and N). In total we compiled changes in 31 potential linear B cell epitopes: 15 for S, 2 for M and 14 for N protein, ranging from one amino acid change to displacement of the reading frame from one to two positions (Supplementary Table 3). In the case of peptides presented by HLA-I, we detected alterations in 72 epitopes, 20 for S protein, 3 for protein M and finally 49 for N protein (Supplementary Table 5). Particularly in the epitopes that are presented by HLA-I, we could observe that even one change in the amino acid sequence, either by substitution or displacement of the reading frame, modifies the immunogenicity index, as well as the specificity for the HLA (Supplementary Table 5). For peptides restricted to HLA-II, we identified 57 epitopes among the variants that showed sequence modifications, 36 belong to S protein, 6 belong to M protein, and 15 in the N protein (Supplementary Table 6).

### Identity analysis between SARS-CoV-2 proteins and other human coronaviruses

To extend our epitope analysis to other human coronaviruses, a multiple sequence alignment of the S protein from SARS-CoV-2 and other human coronaviruses was performed. We identified 25 linear B cell epitopes (Figure 4A) (Supplementary Table 8) that shared a percentage of identity between the S protein of SARS-CoV-2 protein and other human coronaviruses (SARS-CoV, MERS-CoV, HKU1, NL63, OC43, and 229E). This analysis allowed the identification of conserved epitopes among these coronaviruses that could be target of cross-reactive antibodies. We identified epitope GQSKRVDFC which was the only preserved among the 6 coronaviruses, with more than 50% identity. The same analysis was performed for M and N proteins. For M protein, 3 epitopes were identified (Supplementary Table 8) that share a significant percentage of identity between the M protein of SARS-CoV-2 and M protein of the viruses described above, all preserved epitopes are in the C-terminal domain of the protein (Figure 4B)^62^. In the nucleocapsid protein, 7 epitopes with identity percentage higher than 50% with the other human coronaviruses were identified (Supplementary Table 8), these preserved epitopes are in the N-terminal domain and the RNA-binding domain (Figure 4C)^77^.

**Figure 4.**
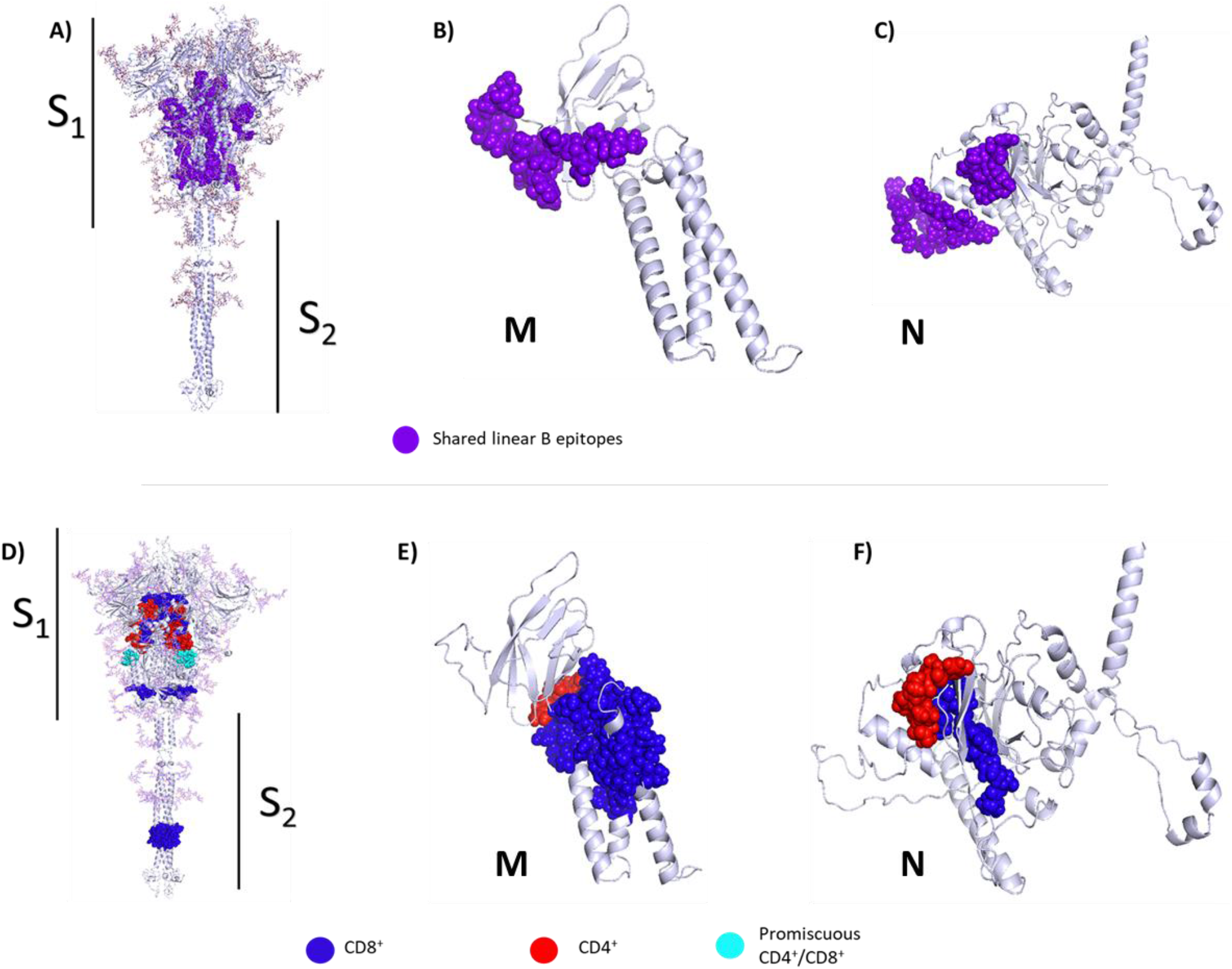
Conserved linear B cell and T cell epitopes in spike, membrane and nucleocapsid proteins of human coronaviruses. 3D graphical representations of the (A, D) spike, (B, E) membrane and (C, F) nucleocapsid proteins of SARS-CoV2. Epitopes that have above 50% identity with epitopes from seasonal coronaviruses (HKU1, NL63, 229E and OC43), SARS-CoV and MERS are highlighted in different colors. Linear B cell epitopes are shown in purple (A, B and C). CD8+ T cell epitopes are represented in blue, CD4+ T cell epitopes are shown in red, and promiscuous epitopes can be seen in cyan (D, E and F). All epitopes (ERIS, EPITS and EEC) were included in the analysis. X-ray crystallography S protein ID: 6VSB_1_1, M protein ID: QHD43419 and N protein ID: QHD43423.

The strategy described above was also used to determine conserved CD8+ or /and CD4+ T cells epitopes. For S protein, 10 different MHC-I-restricted epitopes (Figure 4 in blue) that share identity among the seven coronaviruses were identified (Supplementary Table 9). Most conserved epitopes are located in fusion peptides (FP), central helical region (CH), C-terminal domain (CTD), and finally in heptad repeat 2 (HR2) protein regions (Figure 4D). Supplementary Table 8 shows the conserved peptide sequences with more than 50% identity between the SARS-CoV-2 sequence and the other six coronaviruses. For epitopes restricted to MHC-II (red), we identified 9 conserved epitopes, following the same distribution pattern as the one described of MHC-I-restricted epitopes. For M protein, we identified, one epitope restricted to MHC-I (blue) and one restricted to MHC-II (red), which shared identity between SARS-CoV-2 and the other 6 human coronaviruses (Figure 4E) (Supplementary Table 9). In the case of N protein, 3 conserved MHC-I epitopes (blue) that shared above 50% identity with other human coronaviruses were identified; these epitopes were located between the RNA-binding domain and the binding site. For MHC-II restricted epitopes (red) only one epitope in the N protein was identified (Supplementary Table 9; Figure 4F). As expected, epitopes between SARS-CoV and SARS-CoV-2 shared the highest identity percentages among coronaviruses. In the case of epitopes that are considered as promiscuous (that can be recognized by both CD8+ and CD4+ T cells) 14 epitopes were found in S protein, of which 2 (cyan) have more than 50% identity. For M protein, 6 promiscuous epitopes were identified, while for the N protein, 7 promiscuous epitopes were found (Supplementary Table 9).

## Discussion

SARS-CoV-2 S protein interacts with the host cells receptor ACE2 to mediate virus infection, for this reason, S protein or its RBD domain have been used to develop most of the anti-COVID-19 vaccines that are currently in use around the globe. During the pandemic progression, several mutations within this protein have been identified in different SARS-CoV-2 variants of concern. At different levels, these mutations render virus resistant to neutralization mediated by antibodies induced following previous infections or vaccination, in addition to a reduced immunity against infection with these variants. Nevertheless, the level of protection against serious disease, hospitalisation and dead has been maintained^78,79^.

Through the literature review and epitope prediction exercises carried out in this work, we were able to compile 116 linear B cell epitopes within S protein, reporting 7 new epitopes, most of them located at: RBD, junction of S1/S2 and at the furin site (Figure 1A). Recent studies identified neutralizing antibodies directed to these sites, in that sense, the new epitopes reported here are potential candidates as targets for neutralizing antibodies^57,80^. Several regions of the S protein have been shown as targets for neutralizing antibodies, among them, RBD is one of the important binding sites. (Figure 1A). It has been reported that the epitope GDEVRQIAPGQTGKIADYNYKLPDD, which overlaps in one of the epitopes predicted in this work (Supplementary Table 1), generates neutralizing antibodies in mice^81^. In addition, other studies have shown that different monoclonal antibodies that interact with multiple residues at this site exhibit neutralizing activity^73,82^. The epitopes ASYQTQTNSPRRARSVASQ and IIAYTMSLGAENSVAYSNN (Supplementary Table 1), predicted in our study, have also been described by Polyiam and collaborators, within the polypeptide (SYQTQTNSPRRARSVASQSIIAYTMSLGAENSVAYSN). These epitopes are located at the incision site of the S1/S2 (Figure 1A), where the RRAR sequence is recognized and cleaved by the furin protease, resulting in the separation of the S1 and S2 domains during virus assembly. Antibodies directed to this immunodominant region could block the cleavage of the S protein during viral invasion process^57^. Alpha and Beta SARS-CoV-2 variants exhibit amino acid substitutions in these epitopes, we could also identify these changes in sequences from virus isolated in several countries such as Estonia, South Africa and Singapore (Supplementary Table 5). Of particular importance, the beta variant presents a change in the IIAYTMSLGAENSVAYSNN peptide, in which the substitution A701V affects this epitope at the cleavage site of the cathepsin L^83^.

The peptides ILPDPSKPSKRS, FIEDLLFNKVTLADAGFFIKQYGDCLG and PSKPSKRSFIEDLLFNKV are neutralizing epitopes located at the S2 excision site (residues 805-842), antibody binding at this region results in the inhibition of the molecule excision. Additionally, we located an epitope at the cytoplasmic site of the S protein (CKFDEDDSEPVLKGVKLHYT-1234-1273) (Figure 1A) (Supplementary Table 1), which has been identified as an immunodominant epitope^71,84^. This epitope has not been identified to generate neutralizing antibodies against the SARS-CoV-2 virus, however, a study with the porcine epidemic diarrhea virus (PEDV) (an alpha coronavirus) reported B cell epitopes for neutralizing antibodies located at the cytoplasmic region of the S protein^85^. These results suggest that this region could be another import target for neutralizing antibodies within SARS-CoV-2 S protein.

Recent studies have reported individuals who recovered from SARS-CoV infection that have neutralizing antibodies against the virus, but are not able to neutralize SARS-CoV-2. However, when these subjects were vaccinated with one or two doses of BNT162b2 mRNA vaccine, they produced high-level, broad-spectrum antibodies responses that could neutralize efficiently all SARS-CoV-2 variants of interest and seasonal human coronaviruses^86,87^. Given that the NTD and RBD are the main targets for neutralization, we searched for epitopes conserved between SARS-CoV and SARS-CoV-2 in these regions that could explain cross-neutralization^88^. We could not find any conserved linear B cell epitopes that had at least 50% identity between the two coronaviruses, suggesting that neutralization at these sites is mainly mediated by conformational epitopes. The main limitation of this study is that the B cell epitopes prediction tools we used does not predict conformational epitopes.

As well as the case of S protein, immunodominant epitopes have also been reported for SARS-CoV-2 M and N proteins. In patients with severe disease treated at the Intensive Care Unit (ICU), it has been found high IgG titers specific to S protein linear B cell epitopes TESNKKFLPFQQFGRDIA, PSKPSKRSFIEDLLFNK and N protein epitope NNAAIVLQLPQGTTLPKG^59^. Additionally, the N protein epitope mentioned above, has been associated with lymphopenia in patients with COVID-19^89^. These S and N epitopes have a low mutation rate (<2%) and could be used as markers for COVID-19 induced immunopathology^89^.

Even though neutralizing antibody responses are involved in protection against COVID-19 induced by SARS-CoV-2 infection or vaccination, T cell immune responses have been identified as an extremely important component of immunity against COVID-19. A study in patients with mild and severe COVID-19 showed the presence of effector and central memory SARS-CoV-2-specific T cells, in particular, mild cases generated higher frequencies of cytokine-producing CD8+ T cells^90^. Strong memory-specific T cell responses to SARS-CoV-2 have also been detected in individuals who had mild and asymptomatic infections, in some cases in the absence of antibody responses^91^. Another study reported T cell responses specific to SARS-CoV-2 peptide stimulation in pre-pandemic samples, which suggest T cell cross-reactivity with seasonal coronaviruses^36^. Although pending of experimental confirmation, our work provides a panel of 33 T cell epitopes that could potentially be involved in cross-reactive T cell responses to different coronaviruses (Supplementary Table 9).

*In vitro* studies have characterized that specific CD4+ and CD8+ lymphocytes of convalescent patients are activated by S protein peptides^92–94^. In addition, a study with BNT162b2 vaccine reported that before vaccination, SARS-CoV-2 S protein specific T cells were found in some individuals, suggesting these clones were induced by seasonal coronaviruses^91^. The presence of cross-reactive T cells has been associated with a better humoral and cellular responses to vaccination, authors also reported the presence of Th1 CD4+ and polyclonal CD8+ T cells as well as central memory CD4+ and CD8+ T cells for 6 months after immunization, suggesting long-lasting T cell responses induced by vaccination^95^. These cross-reactive epitopes are located in the S2 region, which, compared to S1 and RBD in particular, is less polymorphic (Supplementary Table 9), supporting the use of these epitopes for the design of broad spectrum vaccines.

In our study, we were able to detect several CD4 and CD8 T cell epitopes, as well as promiscuous epitopes, most of which are located in the S2 region (Supplementary Table 2), the study of epitopes of this region could be also relevant to study the memory T cell compartment induced by SARS-CoV-2 infection or vaccination. In addition of the S protein epitopes reported here, we found 83 and 105 epitopes within M and N proteins, respectively. In the case of M protein only one MHC-I and one MHC-II epitope were found; whereas for N protein, only two MHC-I were found (Figure 2B and 2C). In a previous study, 34 participants with severe and mild COVID-19 responded to M protein peptides, 11 responded to GAVILRLRGHLRIAGHHLGR, 16 to the TSRTLSYYKLGASQRVA, and 3 to the LLESELVIGAVILRGHLR (Supplementary Table 2). The first two activated CD4+ T cells and the later CD8+ T cells ^63^. These peptides are also found by our analysis. Furthermore, the M protein epitope LRGHLRIAGHHLGRCDIKDL, has been described as a highly conserved epitope. A study conducted by Heide and collaborators reported that this peptide was recognized by 12 out of 34 patients with COVID-19, inducing CD4+ T cells to polarize to effector memory phenotype. These data suggest that M protein epitopes could be also relevant in the induction of immunity against SARS-CoV-2^96^.

As for N protein, a study reported that in 19 out of 37 donors of peripheral blood cells who had not been exposed to SARS-CoV nor SARS-CoV-2, the presence of SARS-CoV-2-specific CD4+ IFNγ T cells was detected^96^, again suggesting potential cross-reactive responses with seasonal coronaviruses. In addition, T cell responses against NSP7 and NSP13 non-structural proteins have been identified in donors with no previous exposure to SARS-CoV or SARS-CoV-2^90^, whereas donor samples from COVID-19 and SARS recovered patients reacted preferentially to N protein. In another study, the characterization of specific responses to N protein in a donor without prior exposure to SARS-CoV and SARS-CoV-2, identified the CD4+ and CD8+ T cells epitope MKDLSPRWFYYLGTGPEAG, considered as a promiscuous epitope by Peng and collaborators; this epitope was also recognized by 12 out of 34 patients who recovered from mild and severe COVID-19^63^. Part of this epitope from position 104 to 113 (Supplementary Table 2), was also found by our analysis, it’s located within an N protein region with a high degree of similitude with MERS-CoV, OC43, and HKU1 N proteins, thus we consider this epitope could be relevant for broadly protective vaccines.

An important concern, is if immunity developed by infection or vaccination by SARS-CoV-2 Wuhan strain and its S protein respectively, could protect against infection, symptomatic mild or severe disease or dead caused by the variants of concern of this virus. Mapping the amino acid changes within S protein from the variants of concern together with B and T cell epitopes reported here, revealed that RBD, NTD, and S1 incision sites (Figure 3), are the main regions of the protein that presented amino acid deletions and substitutions. These changes are found in B and T cell epitopes and might alter their antigenicity. Such is the case of several B cell epitopes (GDEVRQIAPGQTGKIADYNYKLPDD, YQAGSTPCNGV, and YGFQPTNGVGYQ) and T cells epitopes (NATRFASVYAWNRK, CVADYSVLYNSASFSKCYGVSPTKLN, DLCFTNVYADSFVI, RQIAPGQTGKIA, TPCNGVEGFNCY, LQSYGFQPTNGVG, and YGYQPYRVVLSF) reported in this study. Additionally, epitope DPFLGVYYHKNNKSWMESEFRVYSSANNCTFEYVSQPFLM is recognized by the monoclonal antibody 4A8^97^, where a deletion in the alpha variant Y144/14 and a substitution in the gamma variant R190S have also been reported (Supplementary Table 5).

The limited information about Omicron variant has revealed that the N440K, G446S, G496S, and Q498R mutations confer the ability to escape antibody responses. It was also found that Q498S and N501S mutations are involved in the immune evasion mechanisms through improving the binding to the ACE2 receptor^98^ (Figure 3) (Supplementary Table 5-7). This might have an impact on vaccines effectiveness, since it has been reported that the sera from patients recovered from infection with the Wuhan strain or vaccinated, present a reduction on neutralizing antibody titres against the variants of concern. The neutralizing capacity of antibodies against Omicron is importantly reduced in healthy and COVID-19 convalescent individuals, who completed their vaccination scheme with BNT162b2 (Pfizer/BioNTech) or mRNA-1273 (Moderna-mRNA) vaccines as well in CoVID-19 convalescent individuals (not vaccinated). This suggests that, despite the vaccination status, protection could be also affected, in particular epidemiological data shows that previously acquired immunity is not highly protective against symptomatic infection with other variants ^99–101^. However, we found multiple cross-reactive B and T cell epitopes that are not modified within the different variants, suggesting that protection against symptomatic or severe disease could be maintained in these individuals despite the protection against infection is compromised. It has been observed patients with a third vaccine dose did not present severe symptoms against different variants including Omicron, which suggests that booster doses amplify the neutralizing capacity of antibodies. In addition, we hypothesize that conserved epitopes could be importantly involved in protection against severe disease or death with novel variants^99,101^.

In conclusion, we found new B and T cell potential epitopes in SARS-CoV-2 S, M and N proteins, and we identified T and B cell epitopes with high percentage of identity with other human coronaviruses. We observe that T cell epitopes have higher identity percentages compared to B cell epitopes. In addition, we mapped the mutations that are within the S protein of SARS-CoV-2 and its variants and observed that most epitopes changes are located on the S1 region. Furthermore, we found that the numbers of changes located within epitopes are distributed in the following order: Delta<Beta<Gamma<Omicron<Alpha. Nevertheless, important number of epitopes remained unchanged among these variants, suggesting that these conserved S protein, M and N proteins epitopes, could mediate cross-protection induced by infection and might be involved in the protection against new SARS-CoV-2 variants induced by Wuhan S protein present in current vaccines. Taken together, this knowledge could be useful for the rational design of novel and broad-spectrum SARS-CoV-2 vaccines.

## Contributors

TR-H and CL-M conceived and designed the study and contributed to data analysis and writing manuscript. SSR-H, DP-O MGG-V contributed to data acquisition, analysis, interpretation and writing manuscript. All authors reviewed the final version. CL-M reviewed and approved the final version.

## Acknowledgments

We would like to thank Guillermo Ramón Torres for kindly facilitating the use of PyMOL® software in order to build the 3D models presented in this manuscript. And to María Guadalupe García Valeriano bachelor’s degree for contributed to data acquisition, analysis, interpretation and witing manuscript.

