## Supplementary Tables 1-9 for "Bioinformatic analysis of B and T cell epitopes from SARS-CoV-2 Spike, Membrane and Nucleocapsid proteins as a strategy to assess possible cross-reactivity between emerging variants, including Omicron, and other human coronaviruses"

### *Supplementary Material*

#### 1 Supplementary Tables

**Supplementary Table 1.** SARS-CoV-2 linear B cell epitopes

| Protein | Sequence | Position | Source |
|---|---|---|---|
| S | MFVFLVLLPLVSSQCVNLTTRTQLPPAYTNSFTRGVY | 1-37 | (1–4) |
| | RSSVLHSTQD | 44-53 | (2,4) |
| | FSNVTWFHAIHV**SGTNGTKRFDN** | 59-81 | (1,2) This study |
| | VYFASTEKSNII | 90-101 | (2) |
| | GTTLDSKTQSLLIVNNATNVVIKVC | 107-131 | (2) |
| | **DPFLGVYYHKNN**KSWMESEFRVYSSANNCTFEYVSQPFLM | 138-177 | (2) This study |
| | MDLEGKQGNFKNL | 177-189 | This study |
| | **KHTPINL**VRDLPQGFS | 203-221 | (2)This study |
| | DLPQGFSALEPLVDLPIGINITRFQTLLALH | 215-245 | (2,5) |
| | RSYLTPGDSSSGWTAGAAAYLLKYNENGTITD | 246-287 | (2,3,5) |
| | YVGYLQPRTFLLKYN | 266-280 | (3) |
| | NGTITD | 282-287 | This study |
| | DAVDCALDPLSETKCTLKSFTVEKGIYQTSN | 287-317 | (2,6) |
| | KSFTVEKGIYQTSNFRVQPTESVRFPNITNLCPFGEVFNATRFASVYAWNRKRISNCVA | 304-363 | (2,3,5,6) |
| | **YNSASFSTFKCYGVSPTKLNDLCFT** | 369-393 | (1,5) |
| | NVYADSFVIR | 394-403 | (5) |
| | **GDEVRQIAPGQTGKIADYNYK**LPDD | 404-426 | (4) This study |
| | NYLYRLFRKSNLKPFERDIS | 450-469 | (4,5) |
| | YQAGSTP**CNGV** | 473-483 | (5)This study |
| | VEGFNCYFPLQ | 483-493 | (3,4) |
| | **YGFQPTNGVGYQ** | 495-506 | (3,4)This study |
| | PYRVVVLSF**ELLHAPATVCGPKKSTNLVKN** | 507-536 | (3,5)This study |
| | KCVNFNFNGLTGTGVLTE | 537-554 | (5) |
| | **SNKKFLPF** | 555-562 | (3,5,7)This study |
| | QQFGRD | 563-568 | (5–7) |
| | **RDIADTTDAVRDPQ** | 567-580 | (5)This study |
| | TLEILDITPCSFGGVSVITPGTNTSNQVAVLYQDV | 581-615 | (5) |

| | | | |
|---|---|---|---|
| | **NCTEVPVAIHADQLTPT** | 616-632 | (5)This study |
| | RVYSTGSNVFQ | 634-644 | This study |
| | VNNSYECDIPI | 656-666 | This study |
| | GAGICASY | 667-674 | (6) |
| | **ASYQTQTNSPRRARSVASQ** | 672-690 | (4)This study |
| | **IIAYTMSLGAENSVAYSN**N | 692-710 | (3,4)This study |
| | GSFCTQLN | 757-764 | (6) |
| | VEQDKNTQE | 772-780 | This study |
| | **KQIYKTPPIKDFGGF** | 786-800 | (4) This study |
| | **ILPDPSKPSKRS** | 805-816 | (4)This study |
| | **FIEDLLFNKVTLADAGFFIKQYGDCLG** | 817-842 | (4,6,7) This study |
| | GAALQIPFAMQMAYRFNGIGVTQNVLYENQKLIANQ | 891-926 | (5,6) |
| | IQDSLSSTASALGKL | 934-948 | This study |
| | DVVNQNAQALNTLVKQLSSNFGAI | 950-973 | (6) |
| | AISSVLNDILSRLDKVEAEVQIDRLITGRLQSLQTYVTQQLIRAAEIRASANLAAT | 972-1027 | (3,6) |
| | **RASANLAATK**MSECVLGQSKRVDFC | 1019-1043 | (6)This study |
| | GYHLM**SFPQSAPHGVVFLHVTY**VPAQEKNFTT | 1056-1077 | (3) This study |
| | RNFYEPQIITTD | 1107-1118 | This study |
| | DVVIGIVNNTVYDPLQPELDSFKEELDKYF**KNHTSPDVDLG**DISGIASVVNIQK | 1124-1181 | (3,6) This study |
| | EIDRLNEVAKNLNESLIDLQELGKYEQY | 1182-1209 | (6) |
| | CKFDEDDSEPVLKGVKLHYT | 1254-1273 | (4,6) |
| M | MADSNGTITVEELKKLLEQWNLVIGFLFLT | 1–30 | (3,4,6,8) |
| | PVTLACFVLAAVYR | 59-72 | (9) |
| | GGIAIAMACLVGLM | 78-91 | (9) |
| | **PLLES**ELVIGAVILRGHLRI | 133–151 | (8) This study |
| | GRCDIKDLPKEITVATSRTLSYYKLGASQRV | 157-187 | (6) |
| | KLGASQRVAGDSGFA | 180-191 | (3) |
| | **YRIGNYKLNTDHSSSSDNIA** | 199-218 | (4)This study |
| | LNTDHSSSSD | 206-215 | (4)This study |
| N | **NGPQNQRNAPRITFGGPSDSTGSNQNGERSGARSKQRRPQGLPNNTASWFTALTQHGK** | 4-61 | (3,6,8) This study |

| | | | |
|---|---|---|---|
| | **QHGKEDLKFPRGQGVPINTNSSPDDQIGYYRRATRRIRGGDGKMKDLS** | 58-105 | (3) This study |
| | **TGPEAGLPYGANK** | 115-127 | (10) This study |
| | GALNTPKDHIGTRNPANNAAIVLQLPQGTTLPKGFYAEGSRGGSQASSRSSSRSRNSSRNSTPG SSRGTSPARMAGNGGD | 137-216 | (3,6,8,10) |
| | LNQLESKMSGKG**QQQQGQTVTKKSAA**EASKKPRQKRTATKAYN | 227-269 | (3,6) This study |
| | YNVTQAFGRRGPEQTQGNF**GDQELIRQGTDYKHWPQIAQFAPSASAF**FGMSRIGMEVTPSGT WL | 268-331 | (3,6)This study |
| | KLDDKDPNFKD | 338-347 | This study |
| | LNKHIDAYKTFPPTEPKKDKKKKADETQALPQRQKKQQTVTLLPAADLDD | 353-403 | (3,6,8) |
| | SKQLQQSMSSADS | 404-416 | This study |

**-In bold the epitopes of our predictions that coincided with the epitopes reported *in silico* (ER)**

**-In red the epitopes with experimental demonstration (ED)**

**Supplementary Table 2.** SARS-CoV-2 proteins T cell epitopes

| Protein | Sequence | Position | Source | HLA affinity | MHC binding |
|---|---|---|---|---|---|
| S | MFVFLVLLPLVSSQCVNLT | 1-19 | This study | HLA-DRB1*01:01, HLA-DRB1*04:01, HLA-DPA1*02:02/DPB1*02:02 | II |
| | YTNSFTRGV | 28-36 | This study | HLA-A*02:01 | I |
| | IWLGFIAGL | 37-45 | This study | HLA-A*23:01 | I |
| | GVYFASIEK | 52-60 | This study | HLA-A*03:01, HLA-A*11:01 | I |
| | LHSTQDLFLPFFSNVTWFHVISGTNGT | 48-74 | This study | HLA-A*02:01, HLA-DRB1*01:01, HLA-DPA1*02:01, HLA-DPB1*02:01, HLA-DPA1*02:02, HLA-DPB1*02:02, HLA-DPA1*03:01, HLA-DPB1*23:01 | I, II |
| | RFDNPVLPF | 78-86 | This study | HLA-A*23:01, HLA-A*24:02 | I |
| | **GVYFASTEK**SNI | 89-100 | (1)<br>This study | HLA-A*03:01, HLA-A*11:01, HLA-DRB1*07:01 | I, II |
| | RGWIFG**TTLDSKTQSLL**IVNATNVVI | 102-128 | (8)<br>This study | HLA-DRB1*01:01, HLA-DRB1*04:01, HLA-DRB1*07:01 | II |
| | NVVIKVCEFQFCNDPFL | 125-141 | This study | HLA-A*02:06, HLA-A*02:01, HLA-DPA1*02:02/DPB1*02:02 | I, II |
| | NKSWMESEFRVY | 144-155 | This study | HLA-A*23:01, HLA-B*35:01, HLA-DPA1*02:02, HLA-DPB1*02:02 | I, II |
| | FRVYSSANN**CTFEYVSQPFLMDLE**GKQGN | 157-185 | (8,11)<br>This study | HLA-A*01:01, HLA-A*11:01, HLA-B*35:01, HLA-DRB1*04:01, HLA-DPA1*02:01/DPB1*02:01, HLA-DPA1*02:02/DPB1*02:02, HLA-DPA1*03:01/DPB1*23:01 | I, II |
| | FKNLREVFKNIDGYF | 187-201 | This study | HLA-A*23:01, HLA-DPA1*02:01, HLA-DPB1*02:01, HLA-DPA1*02:02, HLA-DPB1*02:02, HLA-DPA1*03:01, HLA-DPB1*23:01 | I, II |
| | LEPLVDLPI | 223-231 | This study | HLA-B*40:01 | I |
| | PIGINITRFQTLLALHRSYLTPGDSSSGWTAGAA | 230-263 | (8) | HLA-DRB1*01:01, HLA-DRB1*04:01, HLA-DPA1*01:03, HLA-DPB1*03:01, HLA-DPA1*02:01, HLA-DPB1*02:01, HLA-DPA1*02:02, HLA-DPB1*02:02, HLA-DPA1*03:01, HLA-DPB1*23:01 | II |

| | | | | | |
|---|---|---|---|---|---|
| | GWTAGAAAYYVGYLQPRTFLLK | 257-278 | (1,12–15) | HLA-A*02:01, HLA-A*02:06, HLA-B*08:01, HLA-B*15:01, HLA-DRB1*01:01, HLA-DPA1*02:02/DPB1*02:02, HLA-DPA1*02:01/DPB1*02:01, HLA-DPA1*03:01/DPB1*23:01 | I, II |
| | CTLKSFTVEKGIYQTSNFRVQPTESI | 301-326 | (8) | HLA-DRB1*04:01, HLA-DRB1*07:01 | II |
| | KGIYQTSNFRVQPTESIVRFPNITNLCP | 310-337 | This study | HLA-DRB1*01:01/HLA-DRB1*04:01, HLA-DRB1*07:01 | I, II |
| | **VRFPNITNLCPF** | 327-338 | (6) | HLA-C*14:02, HLA-B*27:05, HLA-B*35:01, HLA-B*51:01, HLA-B*53:01, HLA-B*07:02, HLA-B*54:01 | I |
| | GEVFNATRFASVYAWNRKR | 339-357 | This study | HLA-B*35:01, HLA-A*11:01, HLA-A*03:01 HLA-DQA1*05:01/DQB1*03:01 | I, II |
| | C**VADYSVLYNSASFST**FKCYGVSPTKLN | 361-388 | (6,8) This study | HLA-A*01:01, HLA-A*26:01, HLA-A*29:02, HLA-A*30:02, HLA-DRB1*04:01, HLA-DRB1*01:01, HLA-DRB1*07:01, HLA-DQA1*05:01, HLA-DQB1*03:01 HLA-DRB1*07:01, HLA-DR8 | I, II |
| | SVLYNLAPFSTFK | 363-375 | This study | HLA-A*11:01, HLA-A*03:01, HLA-DPA1*02:02, HLA-DPB1*02:02 | I, II |
| | DLCFTNVYADSFVI | 389-402 | This study | HLA-A*02:01, HLA-DRB1*07:01 | I, II |
| | RQIAPGQTGKIA | 408-419 | This study | HLA-B*07:02, HLA-DQA1*05:01/DQB1*03:01 | I, II |
| | KLPDDFTGCV | 424-433 | (8) | HLA-A*02:01 | I |
| | CVIAWNSNK | 429-437 | This study | HLA-A*11:01, HLA-A*03:01 | I |
| | NYNYLYRLFRKSNLKPFERDISTEIYQ | 448-474 | (8) | HLA-A*24:02, HLA-DRB1*04:01 | I, II |
| | **TPCNGVEGF**NCY | 478-489 | (8,12,13) This study | HLA-A*02:01 | I |
| | LQSYGFQPTNGVG | 492-503 | This study | HLA-B*35:01, HLA-DRB1*04:01 | I, II |
| | **YGYQPYRVVVLSFELL** | 503-518 | (6) This study | HLA-A*23:01, HLA-A*24:02, HLA-A*29:02 HLA-B*07:02, HLA-B*53:01 HLA-A*01:01, HLA-A*26:01, HLA-B*08:01, HLA-DRB1*07:01 | I, II |
| | KKSTNLVKNKCV | 528-539 | This study | HLA-DRB1*07:01 | II |
| | **KCVNF**NFNGLTGTGVLTESN | 537-556 | (6) This study | HLA-DRB1*01:01, HLA-DRB1*07:01, HLA-DQA1*05:01/DQB1*03:01 | II |

| | | | | | |
|---|---|---|---|---|---|
| | **SNKKFLP**FQQFGRD | 552-566 | This study | HLA-DPA1*02:02, HLA-DPB1*02:02 | II |
| | **TPCSFGG**VSVITPGTN | 588-603 | (6)<br>This study | HLA-A*03:01, HLA-DRB1*01:01,<br>HLA-DQA1*05:01/DQB1*03:01 | I, II |
| | YQDVNCTEV | 612-620 | This study | HLA-A*02:01 | I |
| | TWRVYSTGSNVFQTRAGCLIGAE | 632-654 | This study | HLA-DRB1*04:01, HLA-DRB1*07:01,<br>HLA-DRB1*07:01,<br>HLA-DQA1*05:01/DQB1*03:01 | II |
| | CDIPIGAGICASYQTQ | 662-677 | (6) | HLA-A*29:02, HLA-A*30:02, HLA-B*35:0,<br>HLA-DQA1*05:01, HLA-DQB1*03:01 | I, II |
| | **QSIIAYTMS**LGAENSVAYSNN | 690-705 | (8)<br>This study | HLA-A*02:01,<br>HLA-DRB1*04:01, HLA-DRB*07:01<br>HLA-DQA1*05:01/DQB1*03:01 | I, II |
| | IPTNFTISVTTEI | 714-726 | This study | HLA-B*35:01, HLA-B*07:02,<br>HLA-DRB1*07:01 | I, II |
| | SVTTEILPV | 721-729 | This study | HLA-A*02:01 | I |
| | TECSNLLLQYGSFCTQLNRALT | 747-768 | (6,11) | HLA-A*02:01, HLA-A*03:01, HLA-A*11:01,<br>HLA-A*31:01, HLA-A*68:01,<br>HLA-DR8, HLA-DRB1*04:01, | I, II |
| | NTQEVFAQV | 774-782 | This study | HLA-A*02:01 | I |
| | VKQIYKTPPIKDFGGF**NFSQILPDPSKS**K | 785-814 | (8,11)<br>This study | HLA-DRB1*01:01, HLA-DRB1*04:01,<br>HLA-DPA1*02:02, HLA-DPB1*02:02 | II |
| | SKRSFIEDLLFNKV**TLADAGFIK** | 813-835 | (6)<br>This study | HLA-A*02:01, HLA-A*02:02, HLA-A*02:03,<br>HLA-A*02:06, HLA-A*68:02, HLA-A*11:01,<br>HLA-A*31:01, HLA-A*33:01, HLA-A*68:01,<br>HLA-A*03:01, HLA-DRB1*01:01 | I, II |
| | AQKFNGLTVLPPLLTDEM | 852-869 | (6,11) | HLA-DRB1*01:01, HLA-DRB1*04:01 | II |
| | **MIAQY**TSALLAGTIT**SGWTFGAGAALQIPFA** | 869-896 | (11)<br>This study | HLA-A*23:01<br>HLA-DRB1*01:01, HLA-DRB1*07:01,<br>HLA-DQA1*05:01/DQB1*03:01<br>HLA-DQA1*01:01/DQB1*05:01 | I, II |

| | | | | | |
|---|---|---|---|---|---|
| | **TSGWTFGAGAALQIPFAM**QMAYRF NGIGVTQNVLYENQK | 883-921 | (6) This study | HLA-A*11:01, HLA-A*68:01, HLA-B*08:01, HLA-DRB1*01:01, HLA-DRB1*04:01, HLA-DRB1*07:01, HLA-DPA1*01:03, HLA-DPB1*03:01, HLA-DPA1*02:02, HLA-DPB1*02:02, HLA-DPA1*02:01, HLA-DPB1*02:01, HLA-DPA1*03:01, HLA-DPB1*23:01 | I, II |
| | **QKLIANQFNSAIGKIQDSL**SSTASALG KL | 920-948 | This study | HLA-A*11:01, HLA-DRB1*04:01, HLA-DQA1*05:01, HLA-DQB1*03:01 | I, II |
| | **ALGKLQDVVNQNAQALNTLVKQLS SNFGAISSVLN**DILSRL | 944-984 | (1,6,8) This study | HLA-A*02:01, HLA-A*11:01, HLA-A*03:01, HLA-A*31:01, HLA-A*68:01, HLA-B*07:02, HLA-DRB1*01:01, HLA-DRB1*04:01, HLA-DQA1*05:01, HLA-DQB1*03:01 | I, II |
| | RLDKVEA**EVQIDRLITGRLQS**LQTYV TQQLIRAAE**IRASANLAATKM**SE | 983-1032 | (6,8,11,15,16) This study | HLA-A*02:01, HLA-A*02:02, HLA-A*02:06, HLA-A*02:03, HLA-A*03:01, HLA-A*11:01, HLA-A*31:01, HLA-A*33:01, HLA-A*68:02, HLA-B*27:05, HLA-B*07:02, HLA-B*40:01, HLA-B*40:02, HLA-B*44:02, HLA-B*44:03, HLA-B*45:01, HLA-DRB1*01:01, HLA-DRB1*04:01, HLA-DQA1*05:01, HLA-DQB1*03:01, DQB1*03:02 | I, II |
| | RVDFCGKGY | 1039-1047 | (6) | HLA-A*30:02, HLA-A*01:01, HLA-A*03:01, HLA-B*15:01, HLA-B*27:05 | I |
| | QS**APHGVVFLHVT**VPAQEK | 1054-1073 | (6) This study | HLA-A*02:01, HLA-A*02:02, HLA-A*02:03, HLA-A*02:06, HLA-A*03:01, HLA-A*11:01, HLA-A*68:02, HLA-B*07:02, HLA-B*54:01, HLA-B*35:01, HLA-B*53:01, HLA-DPA1*02:02, HLA-DPB1*02:02 | I, II |
| | AQEKNFTTAPAICHD | 1070-1084 | This study | HLA-DQA1*05:01, HLA-DQB1*03:01, HLA-DRB1*07:01 | II |
| | FPREGVFVS | 1089-1097 | This study | HLA-B*35:01, HLA-B*07:02 | I |
| | **FVSNGTHWFVTQRNF**YEPQIITTDNT FVSG | 1095-1124 | (8) This study | HLA-B*35:01, HLA-A*11:01, HLA-A*03:01, HLA-A*23:01, HLA-A*24:02, HLA-DRB1*04:01 | I, II |

| | | | | | |
|---|---|---|---|---|---|
| | IITTDNTFV | 1114-1122 | (6) | HLA-A*02:01 | I |
| | VYDPLQPEL | 1137-1145 | (6) | HLA-A*24:02, HLA-A*29:02, HLA-A*30:02 | I |
| | DSFKEELDKY | 1146-1155 | (6) | HLA-A*26:01, HLA-A*29:02, HLA-A*30:02, HLA-A*01:01 | I |
| | LDKYFKNHTS**PDVDLGDISGINASVV**NIQKEIIDRLNEVAKNL | 1152-1193 | (6,8) This study | HLA-A*02:01, HLA-A*11:01, HLA-B*40:01, HLA-DRB1*01:01, HLA-DRB1*04:01 | I, II |
| | **NLNESLIDLQELGKYEQYIKPWYIWLGFI**AGLIAIV | 1192-1228 | (6,8) This study | HLA-A*02:01, HLA-A*03:01, HLA-A*11:01, HLA-A*23:01, HLA-A*31:01, HLA-A*68:01, HLA-A*33:01, HLA-A*30:02, HLA-A*01:01, HLA-A*26:01, HLA-A*29:02, HLA-B*44:02, HLA-B*18:01, HLA-B*44:03, HLA-B*40:01, HLA-B*40:02, HLA-B*45:01 | I, II |
| | WLGFIAGLIAIVMVTI | 1217-1232 | (6) | HLA-A*02:01, HLA-A*02:02, HLA-A*02:03, HLA-A*02:06, HLA-A*68:02, HLA-A*02:02, HLA-A*02:03, HLA-A*02:01, HLA-A*02:06, HLA-A*68:02, HLA-DRB1*01:01, HLA-DQA1*05:01, HLA-DQB1*03:01 | I, II |
| | **CMTSCCSCLK** | 1236-1245 | (6) | HLA-A*68:01, HLA-A*03:01, HLA-A*11:01, HLA-A*31:01, HLA-A*33:01 | I |
| | DDSEPVLKGVKLHYT | 1259-1273 | (1,6) | HLA-B*40:01, HLA-B*40:02, HLA-B*07:02, HLA-DRB1*01:01 | I, II |
| **M** | **GTITVEELK** | 6-14 | (17) This study | HLA-A*11:01, HLA-A*68:01, HLA-A*03:01, HLA-DPA1*02:02/DPB1*02:02 | I, II |
| | **LKKLLEQWNLVIGFLFLTWICLLQFAYANRNRFLYIIKLIFLWLLWPVTLACF** | 13-65 | (3,6,17,18) This study | HLA-A*02:01, HLA-A*02:02, HLA-A*02:03, HLA-A*02:06, HLA-A*23:01, HLA-A*24:02, HLA-A*29:02, HLA-A*26:01, HLA-A*32:01, HLA-A*68:02, HLA-B*44:03, HLA-B*15:01, HLA-B*44:02, HLA-B*40:01, HLA-B*58:01, HLA-B*57:01 HLA-B*18:01, HLA-B*40:02, HLA-B*45:01, HLA-B*35:01, HLA-DRB1*01:01, HLA-DPA1*01:03, HLA-DPB1*05:01, HLA-DPB1*02:01, HLA-DPB1*04:02, HLA-DPB1*09:01, HLA-DQA1*01:01, HLA-DQA1*01:02, HLA-DQA1*01:03, HLA-DQA1*03:01, HLA-DQB1*03:02, HLA-DQB1*05:01, HLA-DQB1*06:01, HLA-DQB1*06:02 | I, II |

| | | | | | |
|---|---|---|---|---|---|
| | **FLWLLWPVTLACFVLAAVYRINWITGGIAIAMACLVGLMWLSYFIASFRLFARTRSMWSFNPETNILLN** | 53-121 | (3,6,8,9,18–20) This study | HLA-A*02:01, HLA-A*02:02, HLA-A*02:03, HLA-A*02:06, HLA-A*23:01, HLA-A*24:02, HLA-A*29:02, HLA-A*30:01, HLA-A*30:02, HLA-A*32:01, HLA-A*68:02 HLA-B*51:01, HLA-B*35:01, HLA-B*39:01, HLA-B*53:01, HLA-B*07:02, HLA-B*54:01, HLA-B*58:01, HLA-B*57:01, HLA-B*27:05 HLA-C*06:02, HLA-C*07:01, HLA-B*14:02, HLA-DRB1*07:01, HLA-DRB1*01:01, HLA-DRB1*01:02, HLA-DRB1*03:05, HLA-DRB1*03:09, HLA-DRB1*04:01, HLA-DRB1*04:05, HLA-DRB1*04:08, HLA-DRB1*04:26, HLA-DRB1*07:01, HLA-DRB1*07:03, HLA-DRB1*08:03, HLA-DRB1*08:13, HLA-DRB1*09:01, HLA-DRB1*11:01, HLA-DRB1*11:02, HLA-DRB1*11:14, HLA-DRB1*11:20, HLA-DRB1*11:21, HLA-DRB1*11:28, HLA-DRB1*13:02, HLA-DRB1*13:05, HLA-DRB1*13:07, HLA-DRB1*13:22, HLA-DRB1*13:23, HLA-DRB1*15:01, HLA-DRB1*15:02, HLA-DRB1*15:06, HLA-DQA1*02:01, HLA-DQA1*03:01, HLA-DQB1*03:01, HLA-DQB1*03:02, HLA-DQB1*03:03, HLA-DQB1*04:01, HLA-DPA1*02:02/DPB1*02:02, HLA-DPA1*02:01/DPB1*02:01, HLA-DPA1*03:01/DPB1*23:01 | I, II |
| | SELVIGAVILR | 136-146 | (6,11) | HLA-B*40:01, HLA-B*40:02, HLA-B*44:03, HLA-B*45:01, HLA-B*18:01, HLA-B*44:02, HLA-A*68:01, HLA-A*33:01, HLA-A*31:01, HLA-A*11:01, HLA-A*03:01 | I |
| | **LRGHLRIAGHHLGRC** | 145-159 | (3,11) This study | HLA-DRB1*01:01, HLA-DRB1*07:01 | II |
| | RCIKDLPKEITV**ATSRTLSYYKLGASQRVAG** | 159-189 | (3,6,9,11) This study | HLA-A*01:01, HLA-A*30:02, HLA-A*26:01, HLA-A*29:02, HLA-B*57:01 HLA-A*03:01, HLA-A*11:01, HLA-A*31:01, HLA-A*33:01, HLA-A*68:01, HLA-DRB1*01:01, HLA-DRB1*07:01, HLA-DRB1*04:01 | I, II |
| | R**VAGDSGFAAYSRY** | 186-199 | (17) This study | HLA-A*26:01, HLA-A*01:01, HLA-A*30:02, HLA-DRB1*01:01, | I, II |

| | | | | | |
|---|---|---|---|---|---|
| | | | | HLA-DQA1*05:01/DQB1*03:01 | |
| | **RYRIG**NYKL | 198-206 | (6)<br>This study | HLA-A*23:01, HLA-A*24:02, HLA-A*30:02,<br>HLA-DQA1*05:01/DQB1*03:01 | I, II |
| | SSDNIALLV | 213-221 | This study | HLA-A*01:01 | I |
| | PRITFGGPSDSTGSN | 13-27 | This study | HLA-DQA1*05:01/DQB1*03:01 | II |
| N | LPNNTASWF | 45-53 | This study | HLA-B*35:01, HLA-B*07:02 | I |
| | FPRGQGVPI | 66-74 | (3,6) | HLA-B*07:02, HLA-B*54:01, HLA-B*08:01,<br>HLA-B*35:01, HLA-B*51:01, HLA-A*02:01,<br>HLA-B*53:01 | I |
| | PDDQIGYYRRATR**RIRGGDGKM** | 80-101 | (18,21)<br>This study | HLA-B*07:02<br>HLA-DPA1*01:03, HLA-DPB1*02:01,<br>HLA-DPB1*04:02, HLA-DPB1*05:01,<br>HLA-DPB1*09:01 | I, II |
| | **LSPRWYFYYLGTGPEAGLP**YGANKD | 104-128 | (6,11,21) | HLA-A*01:01, HLA-A*29:02, HLA-A*30:02,<br>HLA-A*11:01, HLA-A*23:01, HLA-A*24:02,<br>HLA-A*31:01 HLA-B*07:02, HLA-B*51:01,<br>HLA-B*53:01, HLA-B*54:01,<br>HLA-DRB1*01:01, HLA-DQA1*05:01,<br>HLA-DQB1*03:01 | I, II |
| | GIIWVATEGALNTPKDHI | 129-146 | (3,6,8)<br>This study | HLA-A*11:01, HLA-A*03:01, HLA-A*68:01,<br>HLA-A*31:01, HLA-A*02:01,<br>HLA-DRB1*01:01 | I, II |
| | GTRNPAN**NAAIVLQLPQGTT**LPKGFYA | 147-173 | (6,8,18)<br>This study | HLA-A*02:01, HLA-A*03:01, HLA-A*11:01,<br>HLA-A*68:01, HLA-A*30:02, HLA-A*29:02,<br>HLA-A*26:01,<br>HLA-DRB1*01:01, HLA-DPA1*02:01,<br>HLA-DPA1*02:02, HLA-DPB1*05:01,<br>HLA-DPB1*09:01, HLA-DQA1*01:01,<br>HLA-DQA1*01:02, HLA-DQB1*06:02,<br>HLA-DQA1*05:01/DQB1*03:01 | I, II |
| | **AEGSRGGSQASSRSSSR** | 173-189 | (6)<br>This study | HLA-A*31:01, HLA-A*68:01, HLA-A*11:01,<br>HLA-A*33:01HLA-B*45:01,<br>HLA-DQA1*05:01, HLA-DQB1*03:01 | I, II |
| | RMAGNGG**DAALALLLLDRLNQ**LESKMSG | 209-235 | (6,8,20)<br>This study | HLA-B*40:01, HLA-A*02:01, HLA-A*03:01,<br>HLA-DRB1*01:01, HLA-DQA1*01:03,<br>HLA-DQA1*05:01, HLA-DQA1*03:01,<br>HLA-DQB1*03:02, HLA-DQB1*03:03,<br>HLA-DQB1*04:01, HLA-DQB1*06:01<br>HLA-DPA1*02:02, HLA-DPB1*02:02,<br>HLA-DPB1*23:01 | I, II |

| | | | | | |
|---|---|---|---|---|---|
| | QQQQGQTVTKKSAAEASKK QK**RTATKAYNVT**QAFGRRG | 239-278 | (6,18) This study | HLA-A*11:01, HLA-A*31:01, HLA-DRB1*01:01, HLA-DRB1*07:01, HLA-DQA1*05:01, HLA-DQB1*03:01 | I, II |
| | KHWPQIAQF | 298-307 | This study | HLA-A*23:01 | I |
| | **WPQIAQFAPSASAFFGMSRIGMEVT PSGTWLTY**TGAIKLDDK | 305-342 | (6,8,20,21) This study | HLA-A*02:02, HLA-A*02:06, HLA-A*02:01, HLA-A*02:03, HLA-A*03:01, HLA-A*11:01, HLA-A*31:01, HLA-A*68:0, HLA-A*33:01, HLA-A*03:01, HLA-B*07:02, HLA-B*35:01, HLA-B*51:01, HLA-B*53:01, HLA-A*01:01, HLA-B*40:01, HLA-A*30:02, HLA-A*26:01, HLA-A*29:02, HLA-B*40:01, HLA-DRB1*01:01, HLA-DRB1*04:05, HLA-DRB1*07:01, HLA-DRB1*08:03, HLA-DQA1*05:01, HLA-DQB1*03:01 | I, II |
| | DPNFKDQVILLNKHIDAY**KTFPPTEPK** KD | 343-371 | (6,8,21,22) This study | HLA-A*24:02, HLA-A*02:01 HLA-A*11:01, HLA-A*03:01, HLA-A*31:01, HLA-A*68:01, HLA-DRB1*01:01, HLA-DRB1*04:01 | I, II |
| | QKKQQTVTLLPAADLDDFS | 386-404 | This study | HLA-B*35:01, HLA-DRB1*09:01, HLA-DRB1*01:01, HLA-DRB1*13:02 | I, II |

**-In bold the epitopes of our predictions that coincided with the epitopes reported *in silico* (ER)**

**-In red the epitopes with experimental demonstration (ED)**

**Supplementary Table 3.** Experimentally reported SARS-CoV-2 proteins epitopes and predicted in this study

| Protein | Position | Cell type | Sequence | Source |
|---|---|---|---|---|
| S | 562-580 | B linear | TESNKKFLPFQQFGRDIA | (7) |
| S | 819-835 | B linear | PSKPSKRSFIEDLLFNKV | (7) |
| S | 21-45 | B linear | TRTQLPPAYTNSFTRGVYYPDKVFRS | (5) |
| S | 221-245 | B linear | SALEPLVDLPIGINITRFQTLLALH | (5) |
| S | 261-285 | B linear | GAAAYYVGYLQPRTFLLKYNENGTI | (5) |
| S | 330-349 | B linear | PNITNLCPFGEVFNATRFAS | (5) |
| S | 375-394 | B linear | STFKCYGVSPTKLNDLCFTN | (5) |
| S | 450-469 | B linear | NYLYRLFRKSNLKPFERDIS | (5) |
| S | 489-499 | B linear | CNGVEGFNCYFPLQSYGFQP | (5) |
| S | 522-646 | B linear | ATVCGPKKSTNLVKNKCVNFNFNGLTGTGVLTESNKKFLPFQQFGRDIA DTTDAVRDPQTLEILDITPCSFGGVSVITPGTNTSNQVAVLYQDVNCTEVP VAIHADQLTPTWRVYSTGSNVFQTR | (5) |
| IDi S | 902-926 | B linear | MAYRFNGIGVTQNVLYENQKLIANQ | (5) |
| RBD S | 369-411 | B linear | YNSASFSTFKCYGVSPTKLNDLCFTNVYADSFVIRGDEVRQIA | (5) |
| S | 16–52 | B linear | VNLTTRTQLPPAYTNSFTRGVYYPDKVFRSSVLHSTQ | (4) |
| RBD S | 439-478 | B linear | NNLDSKVGGNYNYLYRLFRKSNLKPFERDISTEIYQAGST | (4) |
| RBD S | 455-478 | B linear | LFRKSNLKPFERDISTEIYQAGST | (4) |
| RBD S | 483-507 | B linear | VEGFNCYFPLQSYGFQPTNGVGYQP | (4) |
| S | 786-815 | B linear | KQIYKTPPIKDFGGFNFSQILPDPSKPSKR | (4) |
| S | 1238-1270 | B linear | TSCCSCLKGCCSCGSCCKFDEDDSEPVLKGVKL | (4) |
| S | 769-786 | B linear | GIAVEQDKNTQEVFAQVK | (10) |
| M | 5-20 | B linear | NGTITVEELKKLLEQW | (4) |
| M | 198-208 | B linear | RYRIGNYKLNTDHSSSSDNIA | (4) |
| N | 153-170 | B linear | NNAAIVLQLPQGTTLPKG | (10) |
| N | 121-138 | B linear | LPYGANKDGIIWVATEGA | (10) |
| N | 129-146 | B linear | GIIWVATEGALNTPKDHI | (10) |
| N | 137-154 | B linear | GALNTPKDHIGTRNPANN | (10) |
| N | 145-162 | B linear | HIGTRNPANNAAIVLQLP | (10) |
| N | 101-120 | T-Cell CD4 | MKDLSPRWYFYYLGTGPEAG | (21) |
| N | 81-95 | T-Cell CD4 | DDQIGYYRRATRRIR | (21) |

| | | | | |
|---|---|---|---|---|
| N | 321-340 | T-Cell CD4/CD8 | GMEVTPSGTWLTYTGAIKLD | (21) |
| N | 104-121 | T-Cell CD4/CD8 | LSPRWYFYYLGTGPEAGL | (11) |
| S | 166-180 | T-Cell CD4/CD8 | CTFEYVSQPFLMDLE | (11) |
| S | 751-765 | T-Cell CD4 | NLLLQYGSFCTQLNR | (11) |
| S | 801-815 | T-Cell CD4 | NFSQILPDPSKPSKR | (11) |
| S | 866-880 | T-Cell CD4 | TDEMIAQYTSALLA | (11) |
| M | 141-158 | T-Cell CD4 | GAVILRGHLRIAGHHLGR | (11) |
| M | 172-188 | T-Cell CD4 | TSRTLSYYKLGASQRVA | (11) |
| M | 133-150 | T-Cell CD8 | LLESELVIGAVILRGHLR | (11) |
| S | 983-1029 | T-Cell CD4 | QLIRAAEIRASANLAATK | (11) |
| S | 269-277 | T- Cell CD8 | YLQPRTFLL | (14,15) |
| S | 1020-1029 | T- Cell CD8 | ASANLAATK | (15,16) |
| S | 257-2654 | T- Cell CD8 | WTAGAAAYY | (12,13) |
| S | 478-486 | T- Cell CD8 | TPCNGVEGF | (12,22) |
| N | 361-369 | T- Cell CD8 | KTFPPTEPK | (12,21) |

**Supplementary Table 4.** Overall description of variants sequences of the SARS-CoV-2 genome

| Name | Tags WHO | ID GISAID | Clade NextClade | CLADE GISAID | LINAGE PANGOLIN |
|---|---|---|---|---|---|
| hCoV-19_Wuhan_NC_045512.2_Ref_NCBI | | | 19A | | B |
| hCoV-19_Saudi/1-29903 Arabia_490010_Clade_GH | | 490010 | 20A | GH | B.1 |
| hCoV-19_Australia_693279_Clade_GV/1-29899 | | 693279 | 20E(EU1) | GV | B.1.177 |
| hCoV-19_Germany_450204_Clade_G/1-29782 | | 450204 | 19A | G | B.1 |
| hCoV-19_Italy_412974_Clade_V/1-29903 | | 412974 | 19A | V | B |
| hCoV-19_Mexico_412972_Clade_GR/1-29903 | | 412972 | 20B | GR | B.1.1.29 |
| hCoV-19_Mexico_K4_1096141_Clade_O/1-29896 | | 1096141 | 20A | O | B.1 |
| hCoV-19_Mexico_K2_1096985_Clade_O/1-29814 | | 1096985 | 20B | O | B.1.1.161 |
| hCoV-19_Taiwan_406031_Clade_O/1-29878 | | 406031 | 19A | O | B |
| hCoV-19_Vietnam_408668_Clade_S/1-29889 | | 408668 | 19A | S | A |
| hCoV-19_Wuhan_Hu-1_402125_Clade_L/1-29903 | | 402125 | 19A | L | B |
| hCoV-19_Wuhan_WH01_406798_Clade_L/1-29866 | | 406798 | 19A | L | B |
| hCoV-19_Australia_NSW3666_710513_Clade_GH/1-29783 | | 710513 | 20C | GH | B.1.517 |
| hCoV-19_Belgium_ULG-11149_737393_Clade_GV/1-29714 | | 737393 | 20E(EU1) | GV | B.1.177 |
| hCoV-19_Ecuador_ZZL-4_660070_Clade_O/1-29903 | | 660070 | 20B | O | B.1.1.4 |
| hCoV-19_Ecuador_USFQ-556_728204_Clade_S/1-29849 | | 728204 | 19B | S | A.28 |
| hCoV-19_Malaysia_718280_Clade_GR/1-29873 | | 718280 | 20B | GR | B.1.1.29 |
| hCoV-19_New Zealand/1-29782 _456378_Clade_V | | 456378 | 19A | V | B.29 |
| hCoV-19_Singapore_728195_Clade_G/1-29879 | | 728195 | 20A | G | B.1.524 |
| hCoV-19_USA_614256_Clade_L/1-29827 | | 614256 | 19A | L | B |
| hCoV-19_Brazil_717944_Clade_GR/1-29864 | Zeta | 717944 | 20B | GR | P.2 |

| | | | | | |
|---|---|---|---|---|---|
| hCoV-19_Brazil_804814_Clade_GR_P.1/1-29606 | Gamma | 804814 | 20J/501Y.V3 | GR | P.1 |
| hCoV-19_Estonia_1101253_Clade_GRY/1-29725 | Alpha | 1101253 | 20I/501Y.V1 | GRY | B.1.1.7 |
| hCoV-19_Italy_717978_Clade_GRY/1-29879 | Alpha | 717978 | 20I/501Y.V1 | GRY | B.1.1.7 |
| hCoV-19_South/1-29872 Africa_KRISP-K002868_602622_Clade_GR | | 602622 | 20B | GR | B.1.1.117 |
| hCoV-19_South/1-29882 Africa_KRISP-EC-K005321_678570_Clade_GH | Beta | 678570 | 20H/501Y.V2 | GH | B.1.351 |
| hCoV-19_England_728343_Clade_GRY/1-29836 | Alpha | 728343 | 20H/501Y.V1 | GRY | B.1.1.7 |
| hCoV-19/Mexico/CDMX-IMSS__K1_1096983_Clade_O | | 1096983 | 19A | O | B |
| hCoV-19/Mexico/CDMX-IMSS_K9_1096142_clade_O | | 1096142 | 19A | O | B |
| hCoV-19_India_1663516_GK | Delta | 1663516 | 21A | GK | B.1.617.2 |
| hCoV-19/India_1663437_Clade_G | | 1663437 | 20A | G | |
| hCoV-19_Botswana_6640916_GRA | Omicron | 6640916 | 21K | GRA | BA.1+B.1.1.529 |
| hCoV-19_Hong/Kong_6590782_GRA | Omicron | 6590782 | 21K | GRA | BA.1+B.1.1.529 |
| hCoV-19_South/Africa_6647956_GRA | Omicron | 6647956 | 21K | GRA | BA.1+B.1.1.529 |

**-In red the sequences of VOC and VBM**

**Supplementary Table 5.** Mutations in epitopes of B cell linear of the SARS-CoV2 and its clinic variants.

| SARS-CoV-2 | Sequence | Country (ID GISAID) | Identity (%) | Lineage |
|---|---|---|---|---|
| | **SQCVNLTTRTQLPPAYTNSFTRGVY** | | | |
| S | SQCVN**F**T**N**RTQLPSAYTNSFTRGVY | Brazil 804814 | 88 | P.1 (γ) |
| | SQCVNL**R**TRTQLPPAYTNSFTRGVY | India 3533076 | 91.3 | B.1.617.2 (δ) |
| | **FSNVTWFHAIHVSGTNGTKRFDN** | | | |
| | FSNVTWFHAI--SGTNGTKRFDN | England 728343 | 91.3 | B.1.1.7 (α) |
| | FSNVTWFHAI--SGTNGTKRFDN | Estonia 1101253 | 91.3 | B.1.1.7 (α) |
| | FSNVTWFHAI--SGTNGTKRFDN | Italy 717978 | 91.3 | B.1.1.7 (α) |
| | FSNVTWFHAIHVSGTNGTKRF**A**N | South Africa 67857 | 95.7 | B.1.351(β) |
| | FSNVTWFHVI—SGTNGTKRFDN | Botswana 66409160 | 91.3 | B.1.1.529 (o) |
| | FSNVTWFHVI—SGTNGTKRFDN | Hong Kong 6530782 | 91.3 | B.1.1.529 (o) |
| | FSNVTWFHVI--SGTNGTKRFDN | South Africa 6647356 | 91.3 | B.1.1.529 (o) |
| | **DPFLGVYYHKNNKSWME** | | | |
| | DPFLGVY-HKNNKSWME | England 728343 | 94.1 | B.1.1.7 (α) |
| | DPFLGVY-HKNNKSWME | Estonia 1101253 | 94.1 | B.1.1.7 (α) |
| | DPFLGVY-HKNNKSWME | Italy 717978 | 94.1 | B.1.1.7 (α) |
| | **Y**PFLGVYYHKNNKSWME | Brazil 804814 | 94.1 | P.1 (γ) |
| | DPFL**D**VYYHKNNKSWME | India 3533076 | 94.1 | B.1.617.2 (δ) |
| | DPFLD**---**HKNNKSWME | Botswana 66409160 | 73.3 | B.1.1.529 (o) |
| | DPFLD**---**HKNNKSWME | Hong Kong 6530782 | 73.3 | B.1.1.529 (o) |
| | DPFLD**---**HKNNKSWME | South Africa 6647356 | 73.3 | B.1.1.529 (o) |
| | **KHTPINLVRDLPQGFS** | | | |
| | KHT**S**INLVRDLPQGFS | Mexico 1096985 | 93.8 | |
| | KHTPINLVR**G**LPQGFS | South Africa 678570 | 93.8 | B.1.351(β) |
| | KHTPI**I**VR**EPE**DLPQGFS | Botswana 66409160 | 73.3 | B.1.1.529 (o) |
| | KHTPI**I**VR**EPE**DLPQGFS | Hong Kong 6530782 | 73.3 | B.1.1.529 (o) |
| | KHTPI**I**VR**EPE**DLPQGFS | South Africa 6647356 | 73.3 | B.1.1.529 (o) |
| | **GDEVRQIAPGQTGKIADYNYKLP** | | | |
| | GDEVRQIAPGQTG**T**IADYNYKLP | Brazil 804814 | 95.7 | P.1 (γ) |
| | GDEVRQIAPGQTG**N**IADYNYKLP | South Africa 678570 | 95.7 | B.1.351 (β) |
| | GDEVRQIAPGQTG**N**IADYNYKLP | Botswana 66409160 | 95.7 | B.1.1.529 (o) |
| | GDEVRQIAPGQTG**N**IADYNYKLP | Hong Kong 6530782 | 95.7 | B.1.1.529 (o) |
| | GDEVRQIAPGQTG**N**IADYNYKLP | South Africa 6647356 | 95.7 | B.1.1.529 (o) |
| | **NLDSKVGGNYNYLYRLFRKSNLKPFERDISTEIYQAGSTPCNGVEGFNCYFPLQSYGFQPTN** | | | |
| | NLDSKVGGNYNYLYRLFRKSNLKPFERDISTEIYQAGSTPCNGVEGFNCYFPLQSYGFQPT**Y** | England 728343 | 98.4 | B.1.1.7 (α) |
| | NLDSKVGGNYNYLYRLFRKSNLKPFERDISTEIYQAGSTPCNGVEGFNCYFPLQSYGFQPT**Y** | Estonia 1101253 | 98.4 | B.1.1.7 (α) |
| | NLDSKVGGNYNYLYRLFRKSNLKPFERDISTEIYQAGSTPCNGVEGFNCYFPLQSYGFQPT**Y** | Italy 717978 | 98.4 | B.1.1.7 (α) |
| | NLDSKVGGNYNYLYRLFRKSNLKPFERDISTEIYQAGSTPCNGV**K**GFNCYFPLQSYGFQPT**Y** | Brazil 804814 | 96.8 | P.1 (γ) |
| | NLDSKVGGN**F**NYLYRLFRKSNLKPFERDISTEIYQAGSTPCNGVEGFNCYFPLQSYGFQPTN | Mexico 1096985 | 98.4 | |
| | NLDSKVGGNYNYLYRLFRKSNLKPFERDISTEIYQAGSTPCNGV**K**GFNCYFPLQSYGFQPTN | Brazil 717944 | 98.4 | |
| | NLDSKVGGNYNYLYRLFRKSNLKPFERDISTEIYQAGSTPCNGVEGFNCYFPLQSYGFQPT**T** | Australia 710513 | 98.4 | |
| | NLDSKVGGNYNYLYRLFRKSNLKPFERDISTEIYQAGSTPCNGV**K**GFNCYFPLQSYGFQPT**Y** | South Africa 602622 | 96.8 | |
| | NLDSKVGGNYNY**R**YRLFRKSNLKPFERDISTEIYQAGS**K**PCNGVEGFNCYFPLQSYGFQPTN | India 3533076 | 96.8 | B.1.617.2 (δ) |
| | **K**LDSKVSGNYNYLYRLFRKSNLKPFERDISTEIYQAG**NK**PCNGV**A**GFNCYFPL**R**SY**S**FRPT**Y** | Botswana 66409160 | 85.5 | B.1.1.529 (o) |
| | **K**LDSKVSGNYNYLYRLFRKSNLKPFERDISTEIYQAG**NK**PCNGV**A**GFNCYFPL**R**SY**S**FRPT**Y** | Hong Kong 6530782 | 85.5 | B.1.1.529 (o) |
| | **K**LDSKVSGNYNYLYRLFRKSNLKPFERDISTEIYQAG**NK**PCNGV**A**GFNCYFPL**R**SY**S**FRPT**Y** | South Africa 6647356 | 85.5 | B.1.1.529 (o) |
| | **ASYQTQTNSPRRARSVASQ** | | | |
| | ASYQTQTNS**H**RRARSVASQ | England 728343 | 94.7 | B.1.1.7 (α) |
| | ASYQTQTNS**H**RRARSVASQ | Estonia 1101253 | 94.7 | B.1.1.7 (α) |
| | ASYQTQTNS**H**RRARSVASQ | Italy 717978 | 94.7 | B.1.1.7 (α) |
| | ASYQTQTNSPRRARSV**V**SQ | South Africa 602622 | 94.7 | |
| | ASYQTQTNS**R**RRARSVASQ | India 3533076 | 94.7 | B.1.617.2 (δ) |
| | ASYQTQT**K**S**H**RRARSVASQ | Botswana 66409160 | 93.8 | B.1.1.529 (o) |
| | ASYQTQT**K**S**H**RRARSVASQ | Hong Kong 6530782 | 93.8 | B.1.1.529 (o) |
| | ASYQTQT**K**S**H**RRARSVASQ | South Africa 6647356 | 93.8 | B.1.1.529 (o) |
| | **YTMSLGAENSVAYSNN** | | | |
| | YTMSLG**V**ENSVAYSNN | Singapore 728195 | 93.8 | |
| | YTMSLG**V**ENSVAYSNN | South Africa 602622 | 93.8 | |
| | YTMSLG**V**ENSVAYSNN | South Africa 678570 | 93.8 | B.1.351 (β) |
| | **RNFYEPQIITTD** | | | |
| | RNFYEPQIITT**H** | England 728343 | 91.7 | B.1.1.7 (α) |
| | RNFYEPQIITT**H** | Estonia 1101253 | 91.7 | B.1.1.7 (α) |
| | RNFYEPQIITT**H** | Italy 717978 | 91.7 | B.1.1.7 (α |
| | **RTQLPPAYTNS** | | | |
| | RTQLP**S**AYTNS | Brazil 804814 | 90.9 | P.1 (γ) |

| | | | | |
|---|---|---|---|---|
| | **SGTNGTKRFDN** | | | |
| | SGTNGTKRFAN | South Africa 678570 | 90.9 | B.1.351(β) |
| | **VRQIAPGQTGKIAD** | | | |
| | VRQIAPGQTGTIAD | Brazil 804814 | 92.9 | P.1 (γ) |
| | VRQIAPGQTGNIAD | South Africa 678570 | 92.9 | B.1.351 (β) |
| | VRQIAPGQTGNIAD | Botswana 66409160 | 92.2 | B.1.1.529 (o) |
| | VRQIAPGQTGNIAD | Hong Kong 6530782 | 92.2 | B.1.1.529 (o) |
| | VRQIAPGQTGNIAD | South Africa 6647356 | 92.2 | B.1.1.529 (o) |
| | **YGFQPTNGVGYQ** | | | |
| | YGFQPTYGVGYQ | England 728343 | 91.7 | B.1.1.7 (α) |
| | YGFQPTYGVGYQ | Estonia 1101253 | 91.7 | B.1.1.7 (α) |
| | YGFQPTYGVGYQ | Italy 717978 | 91.7 | B.1.1.7 (α) |
| | YGFQPTYGVGYQ | Brazil 804814 | 91.7 | P.1 (γ) |
| | YGFQPTYGVGYQ | Australia 710513 | 91.7 | |
| | YGFQPTYGVGYQ | South Africa 678570 | 91.7 | B.1.351(β) |
| | YSFRPTYGVGHQ | Botswana 66409160 | 66.7 | B.1.1.529 (o) |
| | YSFRPTYGVGHQ | Hong Kong 6530782 | 66.7 | B.1.1.529 (o) |
| | YSFRPTYGVGHQ | South Africa 6647356 | 66.7 | B.1.1.529 (o) |
| | **RDIADTTDAVRDPQ** | | | |
| | RDIDDTTDAVRDPQ | England 728343 | 92.9 | B.1.1.7 (α) |
| | RDIDDTTDAVRDPQ | Estonia 1101253 | 92.9 | B.1.1.7 (α) |
| | RDIDDTTDAVRDPQ | Italy 717978 | 92.9 | B.1.1.7 (α) |
| | **QTQTNSPRRARSV** | | | |
| | QTQTNSHRRARSV | England 728343 | 92.3 | B.1.1.7 (α) |
| | QTQTNSHRRARSV | Estonia 1101253 | 92.3 | B.1.1.7 (α) |
| | QTQTNSHRRARSV | Italy 717978 | 92.3 | B.1.1.7 (α) |
| | QTQTNSRRRARSV | India 3533076 | 92.3 | B.1.617.2 (δ) |
| | QTQTKSHRRARSV | Botswana 66409160 | 84.6 | B.1.1.529 (o) |
| | QTQTKSHRRARSV | Hong Kong 6530782 | 84.6 | B.1.1.529 (o) |
| | QTQTKSHRRARSV | South Africa 6647356 | 84.6 | B.1.1.529 (o) |
| | **YRIGNYKLNTDHSSSSDNIA** | | | |
| M | YRIGNYKLNIDHSSSSDNIA | Australia 710513 | 95 | |
| | **LNTDHSSSSD** | | | |
| | LNIDHSSSSD | Australia 710513 | 90 | |
| | **FGGPSDSTGSNQNGERSGARSKQRRPQGLPNN** | | | |
| | FGGPSDSIGSNQNGERSGARSKQRRPQGLPNN | Mexico 1096985 | 96.9 | |
| **N** | FGGPSDSTGSNQNG---GARSKQRRPQGLPNN | Botswana 66409160 | 90.6 | B.1.1.529 (o) |
| | FGGPSDSTGSNQNG---GARSKQRRPQGLPNN | Hong Kong 6530782 | 90.6 | B.1.1.529 (o) |
| | FGGPSDSTGSNQNG---GARSKQRRPQGLPNN | South Africa 6647356 | 90.6 | B.1.1.529 (o) |
| | **HGKEDLKFPRGQGVPINTNSSPDDQIGYYRRATRRIRGGDGKMKDLS** | | | |
| | HGKEDLKFPRGQGVPINTNSSRDDQIGYYRRATR RIRGGDGKMKDLS | Brazil 804814 | 97.9 | P.1 (γ) |
| | HGKEDLKFPRGQGVPINTNSSP-DQIGYYRRATR RIRGGDGKMKDLS | Mexico 1096141 | 97.9 | |
| | HGKEGLKFPRGQGVPINTNSSPDDQIGYYRRATRRIRGGDGKMKDLS | India 3533076 | 97.9 | B.1.617.2 (δ) |
| | **AGLPYGANK** | | | |
| | SGLPYGANK | Brazil 717944 | 88.9 | |
| | **GALNTPKDHIGTRNPANNAAIVLQLPQ** | | | |
| | GAFNTPKDHIGTRNPANNAAIVLQLPQ | South Africa 602622 | 96.3 | |
| | **TTLPKGFYAEGSRGGSQASSRSSSRSRNSSRNSTPGSSRGTSPARMAGNGGD** | | | |
| | TTLPKGFYAEGSRGGSQASSRSSSRSRNSSRNSTPGSSKRTSPARMAGNGGD | England 728343 | 96.2 | B.1.1.7 (α) |
| | TTLPKGFYAEGSRGGSQASSRSSSRSRNSSRNSTPGSSKRTSPARMAGNGGD | Estonia 1101253 | 96.2 | B.1.1.7 (α) |
| | TTLPKGFYAEGSRGGSQASSRSSSRSRNSSRNSTPGSSKRTSPARMAGNGGD | Italy 717978 | 96.2 | B.1.1.7 (α) |
| | TILPKGFYAEGSRGGSQASSRSSSRSRNSSRNSTPGSSKRTSPARIAGNGGD | Ecuador 660070 | 92.3 | |
| | TTLPKGFYAEGSRGGSQASSRSSSRSRNSSRNSTPGSSKRTSPARMAGNGGD | Brazil 804814 | 96.2 | P.1 (γ) |
| | TTLPKGFYAEGSRGGSQASSRSSSRSRNSSRNSTPGSSRGTSPAIMAGNGGD | Mexico 1096141 | 98.1 | |
| | TTLPKGFYAEGSRGGSQASSRSSSRSRNSSRNSTPGSSRGTFPARMAGNGGD | Australia 693279 | 98.1 | |
| | TTLPKGFYAEGSRGGSQASSRSSSRSRNSSRNSTPGSSKRTSPAIMAGNGGD | Malaysia 718280 | 94.2 | |
| | TTLPKGFYAEGSRGGSQASSRSSSRSRNSSRNSTPGSSKRTSPARMAGNGGD | Mexico 412972 | 96.2 | |
| | TTLPKGFYAEGSRGGSQASSRSSSRSRNSSRNSTPGSSKRTSPARMAGNGGD | Brazil 717944 | 96.2 | |
| | TTLPKGFYAEGSRGGSQASSRSSSRSRNSLRNSTPGSSRGTSPARMAGNGGD | Singapore 728195 | 98.1 | |
| | TTLPKGFYAEGSRGGSQASSRSSSRSRNSSRNSTPGSSKRTSPARMAGNGGD | South Africa 602622 | 96.2 | |
| | TTLPKGFYAEGSRGGSQASSRSSSRSRNSSRNSTPGSSRGISPARMAGNGGD | South Africa 678570 | 98.1 | B.1.351(β) |
| | TTLPKGFYAEGSRGGSQASSRSSSRSRNSSRNSTPGSSMGTSPARMAGNGC | India 3533076 | 94.2 | B.1.617.2 (δ) |
| | TTLPKGFYAEGSRGGSQASSRSSSRSRNSSRNSTPGSSKRTSPARMAGNGG | Botswana 66409160 | 96.2 | B.1.1.529 (o) |
| | TTLPKGFYAEGSRGGSQASSRSSSRSRNSSRNSTPGSSKRTSPARMAGNGG | Hong Kong 6530782 | 96.2 | B.1.1.529 (o) |
| | TTLPKGFYAEGSRGGSQASSRSSSRSRNSSRNSTPGSSKRTSPARMAGNGG | South Africa 6647356 | 96.2 | B.1.1.529 (o) |
| | **RLNQLESKMSGKGQQQQGQTVTKKSAAEASKKPRQKRTATKA** | | | |
| | RLNQLESKMFGKGQQQQGQTVTKKSAAEASKKPRQKRTATKA | England 728343 | 97.6 | B.1.1.7 (α) |
| | RLNQLESKMFGKGQQQQGQTVTKKSAAEASKKPRQKRTATKA | Estonia 1101253 | 97.6 | B.1.1.7 (α) |
| | RLNQLESKMFGKGQQQQGQTVTKKSAAEASKKPRQKRTATKA | Italy 717978 | 97.6 | B.1.1.7 (α) |

| | | | | |
|---|---|---|---|---|
| | RLNQLESK**I**SGKGQQQQGQTVTKKSAAEASKKPRQKRTATKA | Brazil 717944 | 97.6 | |
| | RLNQLESK**I**SGKGQQQQGQTVTKKSAAEASKKPRQKRTATKA | Ecuador 660070 | 97.6 | |
| | **DAYKTFPPTEPKKDKKKKADETQALPQRQKKQQTVTLLPAADLDD** | | | |
| | DAYKTFP**S**TEPKKDKKKKADETQALPQRQKKQQTVTLLPAADLDD | Belgium 737393 | 97.8 | |
| | DAYKTFPPTEPKKDKKKKA **Y**ETQALPQRQKKQQTVTLLPAADLDD | India 3533076 | 97.8 | B.1.617.2 (δ) |
| | **QHGKEDLKFPRGQGVPINTNSSPDDQIG** | | | |
| | QHGKEDLKFPRGQGVPINTNSS**R**DDQIG | Brazil 804814 | 96.4 | P.1 (γ) |
| | QHGKEDLKFPRGQGVPINTNSSP**-**DQIG | Mexico 1096141 | 96.4 | |
| | QHGKE**G**LKFPRGQGVPINTNSSPDDQIG | India 3533076 | 96.4 | B.1.617.2 (δ) |
| | **TGPEAGLPYGANK** | | | |
| | TGPE**S**GLPYGANK | Brazil 717944 | 92.3 | |
| | **GALNTPKDHIGTRNPANN** | | | |
| | GA**F**NTPKDHIGTRNPANN | South Africa 602622 | 94.4 | |
| | **GTTLPKGFYAEGSRGGSQASSRSSSRSRNSSRNSTPGSSRGTSPARMAGNGGD** | | | |
| | GT**I**LPKGFYAEGSRGGSQASSRSSSRSRNSSRNSTPGSSRGTSPARMAGNGGD | Ecuador 660070 | 98.1 | |
| | **SKMSGKGQQQQGQTVTKKSAAEASKKPRQKRTATKAYN** | | | |
| | SKM**F**GKGQQQQGQTVTKKSAAEASKKPRQKRTATKAYN | England 728343 | 97.4 | B.1.1.7 (α) |
| | SKM**F**GKGQQQQGQTVTKKSAAEASKKPRQKRTATKAYN | Estonia 1101253 | 97.4 | B.1.1.7 (α) |
| | SKM**F**GKGQQQQGQTVTKKSAAEASKKPRQKRTATKAYN | Italy 717978 | 97.4 | B.1.1.7 (α) |
| | SK**I**SGKGQQQQGQTVTKKSAAEASKKPRQKRTATKAYN | Ecuador 660070 | 97.4 | |
| | SK**I**SGKGQQ QQGQTVTKKSAAEASKKPRQKRTATKAYN | Australia 710513 | 97.4 | |
| | **EVTPSGTWL** | | | |
| | EVT**S**SGTWL | Estonia 1101259 | 88.9 | B.1.1.7 (α) |
| | **KTFPPTEPKKDKKKKADETQALPQRQKKQQ** | | | |
| | KTFP**S**TEPKKDKKKKADETQALPQRQKKQQ | Belgium 737393 | 96.7 | |

**-In red the amino acid changes between the Wuhan sequence and the sequences of other countries.**

**Supplementary Table 6.** Mutations in epitopes of T cell MHC-I of the SARS-CoV2 and its clinic variants.

| SARS-CoV-2 | Sequence | Allele | Country (ID GISAID) | Identity (%) | Immunogenicity score | Lineage |
|---|---|---|---|---|---|---|
| | **YLQPRTFLL** | HLA-A*02:01 | | | 0.1305 | |
| | YLQLRTFLL | | Australia 693279 | 88.9 | 0.1305 | |
| S | **SPRRARSVA** | HLA-B*07:02 | | | | |
| | SHRRARSVA | | Estonia 1101253 | 88.9 | 0.0402 | B.1.1.7 (α) |
| | SHRRARSVA | | Italy 717978 | 88.9 | 0.0402 | B.1.1.7 (α) |
| | SHRRARSVA | | England 728343 | 88.9 | 0.0402 | B.1.1.7 (α) |
| | SPRRARSVV | | South Africa 602622 | 88.9 | 0.0402 | |
| | SRRRARSVA | | India 3533076 | 88.9 | 0.0402 | B.1.617.2 (δ) |
| | SHRRARSVA | | Botswana 6640916 | 88.9 | 0.0402 | B.1.1.529 (o) |
| | SHRRARSVA | | Hong Kong 6530782 | 88.9 | 0.0402 | B.1.1.529 (o) |
| | SHRRARSVA | | South Africa 6647356 | 88.9 | 0.0402 | B.1.1.529 (o) |
| | **ASANLAATK** | HLA-A*11:01 | | | 0.08792 | |
| | ASANLAAIK | HLA-A*03:01 | Brazil 804814 | 88.9 | 0.143 | P.1 (γ) |
| | **NYNYLYRLF** | HLA-A*24:02 | | | 0.0171 | |
| | NFNYLYRLF | HLA-A*23:01 | Mexico 1096985 | 88.9 | 0.0171 | |
| | NYNYRYRLF | | India 3533076 | 88.9 | 0.0171 | B.1.617.2 (δ) |
| | **WTAGAAAYY** | HLA-A*01:01 | | | 0.15259 | |
| | WTAGSAAYY | | Australia 693279 | 88.9 | -0.04661 | |
| | **YQDVNCTEV** | HLA-A*02:01 | | | 0.08295 | |
| | YQGVNCTEV | | Brazil 804814 | 88.9 | 0.08675 | P.1 (γ) |
| | YQGVNCTEV | | South Africa 678570 | 88.9 | 0.08675 | B.1.351(β) |
| | YQGVNCTEV | | Estonia 1101253 | 88.9 | 0.08675 | B.1.1.7 (α) |
| | YQGVNCTEV | | Italy 717978 | 88.9 | 0.08675 | B.1.1.7 (α) |
| | YQGVNCTEV | | England 728343 | 88.9 | 0.08675 | B.1.1.7 (α) |
| | YQGVNCTEV | | Brazil 717944 | 88.9 | 0.08675 | |
| | YQGVNCTEV | | Australia 693279 | 88.9 | 0.08675 | |
| | YQGVNCTEV | | South Africa 602622 | 88.9 | 0.08675 | |
| | YQGVNCTEV | | Singapore 728195 | 88.9 | 0.08675 | |
| | YQGVNCTEV | | Belgium 737393 | 88.9 | 0.08675 | |
| | YQGVNCTEV | | Australia 710513 | 88.9 | 0.08675 | |
| | YQGVNCTEV | | Saudí Arabia 490010 | 88.9 | 0.08675 | |
| | YQGVNCTEV | | Germany 450204 | 88.9 | 0.08675 | |
| | YQGVNCTEV | | Mexico 412972 | 88.9 | 0.08675 | |
| | YQGVNCTEV | | Malaysia 718280 | 88.9 | 0.08675 | |
| | YQGVNCTEV | | India 3533076 | 88.9 | 0.08675 | B.1.617.2 (δ) |
| | YQGVNCTEV | | Botswana 6640916 | 88.9 | 0.08675 | B.1.1.529 (o) |
| | YQGVNCTEV | | Hong Kong 6530782 | 88.9 | 0.08675 | B.1.1.529 (o) |
| | YQGVNCTEV | | South Africa 6647356 | 88.9 | 0.08675 | B.1.1.529 (o) |
| | **VTWFHAIHV** | HLA-A*02:01 | | | 0.38925 | |
| | VTWFHAI-- | | Estonia 1101253 | 77.8 | 0.25803 | B.1.1.7 (α) |
| | VTWFHAI-- | | Italy 717978 | 77.8 | 0.25803 | B.1.1.7 (α) |
| | VTWFHAI— | | England 728343 | 77.8 | 0.25803 | B.1.1.7 (α) |
| | VTWFHAI-- | | Botswana 6640916 | 77.8 | 0.25803 | B.1.1.529 (o) |
| | VTWFHAI-- | | Hong Kong 6530782 | 77.8 | 0.25803 | B.1.1.529 (o) |
| | VTWFHAI-- | | South Africa 6647356 | 77.8 | 0.25803 | B.1.1.529 (o) |
| | **INITRFQTL** | HLA-B*08:01 | | | 0.16778 | |
| | INITRFQT- | | South Africa 678570 | 88.9 | 0.1451 | B.1.351(β) |
| | **IPTNFTISV** | HLA-B*07:02 | | | 0.17229 | |
| | IPINFTISV | HLA-B*35:01 | Estonia 1101253 | 88.9 | 0.20289 | B.1.1.7 (α) |
| | IPINFTISV | | Italy 717978 | 88.9 | 0.20289 | B.1.1.7 (α) |
| | IPINFTISV | | England 728343 | 88.9 | 0.20289 | B.1.1.7 (α) |
| | **LQPRTFLLK** | HLA-A*11:01 | | | 0.18064 | |
| | LQLRTFLLK | | Australia 693279 | 88.9 | 0.18064 | |
| | **FIEDLLFNK** | HLA-A*11:01 | | | 0.1286 | |
| | FIEDLLFDK | | Mexico 1096141 | 77.8 | 0.14534 | |
| | **KNLREFVFK** | HLA-A*11:01 | | | 0.35942 | |

| | | | | | | |
|---|---|---|---|---|---|---|
| | KNLSEFVFK | | Brazil 804814 | 88.9 | 0.14087 | P.1 (γ) |
| | **RFDNPVLPF** | HLA-A*24:02 | | | 0.01291 | |
| | RFANPVLPF | HLA-A*23:01 | South Africa 678570 | 88.9 | 0.01291 | B.1.351(β) |
| | **GAAAYYVGY** | HLA-A*11:01 | | | 0.09963 | |
| | GSAAYYVGY | HLA-B*35:01 | Australia 693279 | 88.9 | 0.09963 | |
| | **YNYLYRLFR** | HLA-A*11:01 | | | 0.0918 | |
| | FNYLYRLFR | | Mexico 1096141 | 88.9 | 0.0918 | |
| | YNYRYRLFR | | India 3533076 | 88.9 | 0.0918 | B.1.617.2 (δ) |
| | **RFDNPVLPF** | HLA-A*23:01 | | | 0.01291 | |
| | RFANPVLPF | HLA-A*24:02 | South Africa 678570 | 88.9 | 0.01291 | B.1.351(β) |
| | **NASVVNIQK** | HLA-A*11:01 | | | 0.06659 | |
| | NASFVNIQK | | Brazil 804814 | 88.9 | 0.14285 | P.1 (γ) |
| | NASFVNIQK | | Brazil 717944 | 88.9 | 0.14285 | |
| | **TPCNGVEGF** | HLA-B*35:01 | | | 0.15215 | |
| | TPCNGVKGF | | Brazil 804814 | 88.9 | -0.11435 | P.1 (γ) |
| | TPCNGVKGF | | South Africa 602622 | 88.9 | -0.11435 | |
| | TPCNGVKGF | | Brazil 717944 | 88.9 | -0.11435 | |
| | TPCNGVKGF | | South Africa 678570 | 88.9 | -0.11435 | B.1.351(β) |
| | KPCNGVEGF | | India 3533076 | 88.9 | 0.15215 | B.1.617.2 (δ) |
| | KPCNGVAGF | | Botswana 6640916 | 77.8 | 0.10067 | B.1.1.529 (o) |
| | KPCNGVAGF | | Hong Kong 6530782 | 77.8 | 0.10067 | B.1.1.529 (o) |
| | KPCNGVAGF | | South Africa 6647356 | 77.8 | 0.10067 | B.1.1.529 (o) |
| | **NGVEGFNCY** | HLA-B*35:01 | | | 0.22039 | |
| | NGVKGFNCY | | Brazil 804814 | 88.9 | -0.09736 | P.1 (γ) |
| | NGVKGFNCY | | South Africa 602622 | 88.9 | -0.09736 | |
| | NGVKGFNCY | | South Africa 678570 | 88.9 | -0.09736 | B.1.351(β) |
| | NGVKGFNCY | | Brazil 717944 | 88.9 | -0.09736 | |
| | NGVAGFNCY | | Botswana 6640916 | 88.9 | 0.15901 | B.1.1.529 (o) |
| | NGVAGFNCY | | Hong Kong 6530782 | 88.9 | 0.15901 | B.1.1.529 (o) |
| | NGVAGFNCY | | South Africa 6647356 | 88.9 | 0.15901 | B.1.1.529 (o) |
| | **QPRTFLLKY** | HLA-B*35:01 | | | 0.02406 | |
| | QLRTFLLKY | | Australia 693279 | 88.9 | 0.02406 | |
| | **ASFSTFKCY** | HLA-A*11:01 | | | -0.19397 | |
| | APFFTFKCY | HLA-B*35:01 | Botswana 6640916 | 77.8 | 0.0903 | B.1.1.529 (o) |
| | **SVLYNSASF** | HLA-B*35:01 | | | -0.23299 | |
| | SVLYNLAPF | | Botswana 6640916 | 77.8 | 0.00248 | B.1.1.529 (o) |
| | SVLYNLAPF | | South Africa 6647356 | 77.8 | 0.00248 | B.1.1.529 (o) |
| | **VNFNFNGLT** | HLA-A*03:01 | | | 0.16152 | |
| | VNFNFNGLK | | Botswana 6640916 | 88.9 | 0.16152 | B.1.1.529 (o) |
| | VNFNFNGLK | HLA-A*11:01 | Hong Kong 6530782 | 88.9 | 0.16152 | B.1.1.529 (o) |
| | VNFNFNGLK | | South Africa 6647356 | 88.9 | 0.16152 | B.1.1.529 (o) |
| | **YNSASFSTF** | HLA-B*35:01 | | | -0,18217 | |
| | YNLAPFFTF | HLA-A*24:02 | Botswana 6640916 | 66.7 | 0.25665 | B.1.1.529 (o) |
| | **YGFQPTNGV** | HLA-B*35:01 | | | -0.03848 | |
| | YGFQPTYGV | | Estonia 1101253 | 87.5 | -0.05828 | B.1.1.7 (α) |
| | YGFQPTYGV | | Italy 717978 | 87.5 | -0.05828 | B.1.1.7 (α) |
| | YGFQPTYGV | | England 728343 | 87.5 | -0.05828 | B.1.1.7 (α) |
| | YGFQPTYGV | | South Africa 678570 | 87.5 | -0.05828 | B.1.351(β) |
| | YGFQPTYGV | | Brazil 804814 | 87.5 | -0.05828 | P.1 (γ) |
| | YSFRPTYGV | HLA-A*02:01 | Botswana 6640916 | 66.7 | 0.1325 | B.1.1.529 (o) |
| | YSFRPTYGV | | Hong Kong 6530782 | 66.7 | 0.1325 | B.1.1.529 (o) |
| | **SVLNDILSR** | HLA-A*31:01 | | | 0.03075 | |
| | SVLNDIFSR | HLA-A*11:01 | Hong Kong 6530782 | 88.9 | 0.13891 | B.1.1.529 (o) |
| | SVLNDIFSR | | South Africa 6647356 | 88.9 | 0.13891 | B.1.1.529 (o) |
| | **VLNDILSRL** | HLA-A*02:01 | | | 0.03 | |
| | VLNDIFSRL | | Hong Kong 6530782 | 88.9 | 0.15064 | B.1.1.529 (o) |
| | VLNDIFSRL | | South Africa 6647356 | 88.9 | 0.15064 | B.1.1.529 (o) |
| M | **KLLEQWNLV** | HLA-A*02:01 | | | 0.18092 | |
| | KLLEEWNLV | | Botswana 6640916 | 88.9 | 0.39122 | B.1.1.529 (o) |
| | KLLEEWNLV | | Hong Kong 6530782 | 88.9 | 0.39122 | B.1.1.529 (o) |
| | KLLEEWNLV | | South Africa 6647356 | 88.9 | 0.39122 | B.1.1.529 (o) |

| | | | | | | |
|---|---|---|---|---|---|---|
| | DLSPRWYIR | | Brazil 804814 | 22.2 | 0.26869 | P.1 (γ) |
| | **VTLACFVLAA** | HLA-A*02:06 | | | 0.14963 | |
| | VTLTCFVLAA | HLA-A*02:01 | Botswana 6640916 | 90 | 0.12471 | B.1.1.529 (o) |
| | VTLTCFVLAA | | Hong Kong 6530782 | 90 | 0.12471 | B.1.1.529 (o) |
| | VTLTCFVLAA | | South Africa 6647356 | 90 | 0.12471 | B.1.1.529 (o) |
| | **TPSGTWLTY** | HLA-B*35:01 | | | 0.24003 | |
| | TSSGTWLTY | | Estonia 1101253 | 88.9 | 0.24003 | B.1.1.7 (α) |
| N | **NGERSGARSK** | HLA-A*02:01 | | | -0.05967 | |
|  | NG----GARSK |  | Botswana 6640916 | 70 | ////* | B.1.1.529 (o) |
|  | NG----GARSK |  | Hong Kong 6530782 | 70 | ////* | B.1.1.529 (o) |
|  | NG----GARSK |  | South Africa 6647356 | 70 | ////* | B.1.1.529 (o) |
| | **KTFPPTEPK** | HLA-A*03:01 | | | 0.1306 | |
| | KTFPSTEPK | HLA-A*11:01 | Belgium 737393 | 88.9 | -0.0197 | |
| | **SPDDQIGYY** | HLA-B*35:01 | | | 0.06844 | |
| | SRDDQIGYY | | Brazil 804814 | 88.9 | 0.06844 | P.1 (γ) |
| | SP-DQIGYY | | Mexico 1096141 | 88.9 | //* | |
| | **AGLPYGANK** | HLA-A*11:01 | | | 0.04278 | |
| | SGLPYGANK | | Brazil 717944 | 88.9 | 0.04278 | |
| | **KHWPQIAQF** | HLA-A*23:01 | | | 0.03856 | |
| | KHWPQIALF | | Mexico 1096141 | 88.9 | 0.09976 | |
| | **FPPTSFGPL** | HLA-B*07:02 | | | 0.00668 | |
| | FPLTSFGP | HLA-B*35:01 | Brazil 804814 | 77.8 | 0.01316 | P.1 (γ) |
| | FPLTSFGP | | Estonia 1101253 | 77.8 | 0.01316 | B.1.1.7 (α) |
| | FPLTSFGP | | Ecuador 660070 | 77.8 | 0.01316 | |
| | FPLTSFGP | | Belgium 737393 | 77.8 | 0.01316 | |
| | FPLTSFGP | | Italy 717978 | 77.8 | 0.01316 | B.1.1.7 (α) |
| | FPLTSFGP | | England 728343 | 77.8 | 0.01316 | B.1.1.7 (α) |
| | FPLTSFGP | | Australia 710513 | 77.8 | 0.01316 | |
| | FPLTSFGP | | Saudi Arabia 490010 | 77.8 | 0.01316 | |
| | FPLTSFGP | | South Africa 602622 | 77.8 | 0.01316 | |
| | FPLTSFGP | | Singapore 728195 | 77.8 | 0.01316 | |
| | FPLTSFGP | | Malaysia 718280 | 77.8 | 0.01316 | |
| | FPLTSFGP | | Brazil 717944 | 77.8 | 0.01316 | |
| | FPLTSFGP | | Mexico 412972 | 77.8 | 0.01316 | |
| | **CSLSHRFYR** | HLA-A*11:01 | | | 0.00679 | |
| | --NSHRFYR | | Australia 693279 | 66.7 | 0.1731 | |
| | **LLTILTSLL** | HLA-A*02:01 | | | 0.02616 | |
| | LITILTSLL | | Australia 710513 | 88.9 | 0.02616 | |
| | **VLFSTVFPP** | HLA-A*02:01 | | | 0.04051 | |
| | VLFSTVFPL | | Brazil 804814 | 88.9 | 0.04051 | P.1 (γ) |
| | VLFSTVFPL | | Estonia 1101253 | 88.9 | 0.04051 | B.1.1.7 (α) |
| | VLFSTVFPL | | Ecuador 660070 | 88.9 | 0.04051 | |
| | VLFSTVFPL | | Belgium 737393 | 88.9 | 0.04051 | |
| | VLFSTVFPL | | Italy 717978 | 88.9 | 0.04051 | B.1.1.7 (α) |
| | VLFSTVFPL | | England 728343 | 88.9 | 0.04051 | B.1.1.7 (α) |
| | VLFSTVFPL | | Australia 710513 | 88.9 | 0.04051 | |
| | VLFSTVFPL | | Saudi Arabia 490010 | 88.9 | 0.04051 | |
| | VLFSTVFPL | | South Africa 602622 | 88.9 | 0.04051 | |
| | VLFSTVFPL | | Singapore 728195 | 88.9 | 0.04051 | |
| | VLFSTVFPL | | Malaysia 718280 | 88.9 | 0.04051 | |
| | VLFSTVFPL | | Brazil 717944 | 88.9 | 0.04051 | |
| | VLFSTVFPL | | Mexico 412972 | 88.9 | 0.04051 | |
| | **CIRCLWSTK** | HLA-A*03:01 | | | 0.04332 | |
| | CIRCLWSIK | | Singapore 728195 | 88.9 | 0.0984 | |
| | **VSALVYDNK** | HLA-A*11:01 | | | 0.0532 | |
| | VSALVYDNR | | Estonia 1101253 | 88.9 | 0.0532 | B.1.1.7 (α) |
| | **MNVLTLVYK** | HLA-A*11:01 | | | 0.06228 | |
| | MNVLTLFYK | | Australia 710513 | 88.9 | 0.12624 | |
| | **IIWFLLLSV** | HLA-A*02:01 | | | 0.06244 | |
| | TIWFLLLSV | | Estonia 1101253 | 88.9 | 0.06244 | B.1.1.7 (α) |
| | TIWFLLLSV | | Italy 717978 | 88.9 | 0.06244 | B.1.1.7 (α) |
| | TIWFLLLSV | | England 728343 | 88.9 | 0.06244 | B.1.1.7 (α) |

| | | | | | | |
|---|---|---|---|---|---|---|
| | IIWFLIILSV | | Wuhan 406798 | 88.9 | 0.19816 | |
| | **GTSTDVVYR**<br>---FDVVYR | HLA-A*11:01 | <br>Germany 450204 | <br>55.6 | 0.0785<br>0.05134 | |
| | **LMNVLTLVY**<br>LMNVLTLFY | HLA-A*01:01<br>HLA-A*03:01 | <br>Australia 710513 | <br>88.9 | 0.07994<br>0.12422 | |
| | **CIDGALLTK**<br>CIDGVLLTK | HLA-A*03:01<br>HLA-A*11:01 | <br>Ecuador 660070 | <br>88.9 | 0.08228<br>0.08438 | |
| | **KQGDDYVYL**<br>KQGDDYLYL | HLA-A*02:01 | <br>Malaysia 718280 | <br>88.6 | 0.08412<br>0.03992 | |
| | **IINNTVYTK**<br>IINNTVYIK | HLA-A*03:01<br>HLA-A*11:01 | <br>Australia 710513 | <br>88.9 | 0.08761<br>0.14269 | |
| | **LLLTILTSL**<br>LLITILTSL | HLA-A*02:01<br>HLA-B*08:01 | <br>Australia 710513 | <br>88.9 | 0.09072<br>0.13752 | |
| | **VLITEGSVK**<br>VLITEGIVK | HLA-A*03:01<br>HLA-A*11:01 | <br>Belgium 737393 | <br>88.9 | 0.09616<br>0.3481 | |
| | **DAVTAYNGY**<br>NAVTAYNGY | HLA-B*35:01 | <br>South Africa 602622 | <br>88.9 | 0.10142<br>0.10142 | |
| | **GEAANFCAL**<br>GEADNFCAL<br>GEADNFCAL<br>GEADNFCAL | HLA-B*40:01<br>HLA-B*44:02<br>HLA-B*44:03 | <br>Estonia 1101253<br>Italy 717978<br>England 728343 | <br>88.9<br>88.9<br>88.9 | 0.13333<br>0.11628<br>0.11628<br>0.11628 | <br>B.1.1.7 (α)<br>B.1.1.7 (α)<br>B.1.1.7 (α) |
| | **FFFLYENAF**<br>FFFFYENAF<br>FFFFYENAF | HLA-A*23:01<br>HLA-B*35:01 | <br>Italy 412974<br>New Zealand 456378 | <br>88.9<br>88.9 | 0.13489<br>0.26385<br>0.26385 | <br>B.1.1.7 (α) |
| | **IFWRNTNPI**<br>IFWRNTNSI | HLA-A*23:01 | <br>Australia 710513 | <br>88.9 | 0.14228<br>0.0521 | |
| | **TLYCIDGAL**<br>TLYCIDGVL | HLA-A*02:01 | <br>Ecuador 660070 | <br>88.9 | 0.14649<br>0.14775 | |
| | **IPRRNVATL**<br>IPRRNVVTL | HLA-B*07:02<br>HLA-B*08:01 | <br>Ecuador 660070 | <br>88.9 | 0.15714<br>0.15896 | |
| | **LYENAFLPF**<br>FYENAFLPF<br>FYENAFLPF | HLA-A*23:01<br>HLA-A*24:02 | <br>Italy 412974<br>New Zealand 456378 | <br>88.9<br>88.9 | 0.15845<br>0.15845<br>0.15845 | <br>B.1.1.7 (α) |
| | **YLITPVHVM**<br>YLIIPVHVM | HLA-A*02:01<br>HLA-B*35:01 | <br>Singapore 728195 | <br>88.9 | 0.16174<br>0.2566 | |
| | **SLSHRFYRL**<br>-NSHRFYRL | HLA-A*02:01<br>HLA-B*08:01 | <br>Australia 693279 | <br>77.8 | 0.16657<br>0.1731 | |
| | **TQWSLFFFL**<br>TQWSLFFFF<br>TQWSLFFFF | HLA-A*02:01<br>HLA-A*23:01 | <br>Italy 412974<br>New Zealand 456378 | <br><br>88.9 | 0.17203<br>0.17203<br>0.17203 | <br>B.1.1.7 (α) |
| | **AIFYLITPV**<br>AIFYLIIPV | HLA-A*02:01 | <br>Singapore 728195 | <br>88.9 | 0.17504<br>0.2546 | |
| | **LYCIDGALL**<br>LYCIDGVLL | HLA-A*23:01<br>HLA-A*24:02 | <br>Ecuador 660070 | <br>88.9 | 0.19646<br>0.19828 | |
| | **SLVPGFNEK**<br>SFVPGFNEK<br>SLVPGFSEK | HLA-A*03:01<br>HLA-A*11:01 | <br>Ecuador 728204<br>Belgium 737393 | <br>88.9<br>88.9 | 0.19848<br>0.19848<br>0.06432 | |
| | **ITVNVLAWL**<br>ITVNVVAWL | HLA-A*02:01 | <br>Brazil 717944 | <br>88.9 | 0.19909<br>0.24839 | |
| | **KVTLVFLFV**<br>KVTLVLLFV | HLA-A*02:01 | <br>Estonia 1101253 | <br>88.9 | 0.21088<br>0.09024 | <br>B.1.1.7 (α) |
| | **FYLITPVHV**<br>FYLIIPVHV | HLA-A*23:01<br>HLA-A*24:02 | <br>Singapore 728195 | <br>88.9 | 0.21142<br>0.30322 | |
| | **FLYENAFLP**<br>FFYENAFLP<br>FFYENAFLP | HLA-A*02:01 | <br>Italy 412974<br>New Zealand 456378 | <br>88.9<br>88.9 | 0.2224<br>0.2224<br>0.2224 | <br>B.1.1.7 (α) |
| | **YFMRFRRAF**<br>YFMRFRRAY | HLA-B*08:01<br>HLA-A*23:01 | <br>Ecuador 728204 | <br>88.9 | 0.22434<br>0.22434 | |
| | **HSIGFDYVY**<br>HSIGLDYVY | HLA-B*35:01 | <br>Ecuador 728204 | <br>88.9 | 0.23318<br>0.10838 | |

| | | | | | | |
|---|---|---|---|---|---|---|
| | **IPARARVEC**<br>IPARARV**D**C | HLA-B*07:02 | <br>Brazil 804814 | <br>88.9 | 0.24494<br>0.1994 | <br>P.1 (γ) |
| | **TVNVLAWLY**<br>TVNV**V**AWLY | HLA-A*01:01<br>HLA-A*11:01 | <br>Brazil 717944 | <br>88.9 | 0.24593<br>0.29693 | |
| | **MEIDFLELA**<br>ME**V**DFLELA | HLA-B*44:03<br>HLA-B*40:01 | <br>Belgium 737393 | <br>88.9 | 0.2471<br>0.2173 | |
| | **NVLAWLYAA**<br>NV**V**AWLYAA | HLA-A*02:01 | <br>Brazil 717944 | <br>88.9 | 0.26077<br>0.27777 | |
| | **TLVFLFVAA**<br>TLV**L**LFVAA | HLA-A*02:01 | <br>Estonia 1101253 | <br>88.9 | 0.2883<br>0.15934 | <br>B.1.1.7 (α) |
| | **NINIVGDFK**<br>**Y**INIVGDFK | HLA-A*11:01 | <br>England 728343 | <br>88.9 | 0.29104<br>0.29104 | <br>B.1.1.7 (α) |
| | **VFLFVAAIF**<br>V**L**LFVAAIF | HLA-A*23:01<br>HLA-A*24:02 | <br>Estonia 1101253 | <br>88.9 | 0.30201<br>0.30201 | <br>B.1.1.7 (α) |
| | **WEPEFYEAM**<br>**C**EPEFYEAM | HLA-B*40:01 | <br>Taiwan 406031 | <br>88.9 | 0.31503<br>0.31503 | |
| | **TPGSSRGTSPAR**<br>TPGSS**KR**TSPAR<br>TPGSS**KR**TSPAR<br>TPGSS**KR**TSPAR<br>TPGSSRG**I**SPAR<br>TPGSS**KR**TSPAR<br>TPGSS**M**GTSPAR<br>TPGSS**KR**TSPAR<br>TPGSS**KR**TSPAR<br>TPGSS**KR**TSPAR | HLA-A*30:02<br>HLA-A*02:01<br><br><br>HLA-A*30:02<br>HLA-A*02:01<br>HLA-A*30:02<br>HLA-A*02:01 | <br>Estonia 1101253<br>Italy 717978<br>England 728343<br>South Africa 678570<br>Brazil 804814<br>India 3533076<br>Botswana 6640916<br>Hong Kong 6530782<br>South Africa 6647356 | <br>83.3<br>83.3<br>83.3<br>91.7<br>83.3<br>91.7<br>83.3<br>83.3<br>83.3 | -0,5806<br>-0,3376<br>-0,3376<br>-0,3376<br>-0.2458<br>-0,3376<br>-0.559<br>-0,3376<br>-0,3376<br>-0,3376 | <br>B.1.1.7 (α)<br>B.1.1.7 (α)<br>B.1.1.7 (α)<br>B.1.351(β)<br>P.1 (γ)<br>B.1.617.2 (δ)<br>B.1.1.529 (o)<br>B.1.1.529 (o)<br>B.1.1.529 (o) |
| | **FMRFRRAFG**<br>FMRFRRA**Y**G | HLA-B*08:01 | <br>Ecuador 728204 | <br>88.9 | 0.33514<br>0.26458 | |
| | **LINIIIWFL**<br>LINI**T**IWFL<br>LINI**T**IWFL<br>LINI**T**IWFL | HLA-A*02:01 | <br>Estonia 1101253<br>Italy 717978<br>England 728343 | <br>88.9<br>88.9<br>88.9 | 0.64204<br>0.55024<br>0.55024<br>0.55024 | <br>B.1.1.7 (α)<br>B.1.1.7 (α)<br>B.1.1.7 (α) |

**-In this data we can not determine the immunogenicity score because it was missing an amino acid (*)**

**-In red the amino acid changes between the Wuhan sequence and the sequences of other countries.**

**Supplementary Table 7.** Mutations in epitopes of T cell MHC-II of the SARS-CoV2 and its clinic variants.

| SARS-CoV-2 | Sequence | Allele | Country (ID GISAID) | Identity (%) | Lineage |
|---|---|---|---|---|---|
| S | **LPLVSSQCVNLT** | HLA-DRB1*04:01 | | | |
| | LPLVSSQCVNFT | | Brazil 804814 | 91.7 | P.1 (γ) |
| | LPLVSSQCVNLR | | India 3533076 | 91.7 | B.1.617.2 (δ) |
| | **TNFTISVTTEIL** | HLA-DRB1*07:01 | | | |
| | INFTISVTTEIL | | Estonia 1101253 | 91.7 | B.1.1.7 (α) |
| | INFTISVTTEIL | | Italy 717978 | 91.7 | B.1.1.7 (α) |
| | INFTISVTTEIL | | England 728343 | 91.7 | B.1.1.7 (α) |
| | **AEIRASANLAAT** | HLA-DRB1*04:01 | | | |
| | AEIRASANLAAI | HLA-DRB1*07:01 | Brazil 804814 | 91.7 | P.1 (γ) |
| | | HLA-DQA1*05:01/DQB1*03:01 | | | |
| | **TRFQTLLALHRS** | HLA-DRB1*01:01 | | | |
| | TRFQT---LHRS | HLA-DRB1*04:01 | South Africa 678570 | 75 | B.1.351(β) |
| | | HLA-DPA1*01:03/DPB1*03:01 | | | |
| | | HLA-DPA1*02:02/DPB1*02:02 | | | |
| | **FIEDLLFNKVTL** | HLA-DPA1*03:01/DPB1*23:01 | | | |
| | FIEDLLFDKVTL | HLA-DPA1*02:02/DPB1*02:02 | | | |
| | | HLA-DPA1*02:01/DPB1*02:01 | Mexico 1096141 | 91.7 | |
| | **QSIIAYTMSLGA** | HLA-DRB1*04:01 | | | |
| | QSIIAYTMSLGV | | South Africa 678570 | 91.7 | B.1.351(β) |
| | QSIIAYTMSLGV | | Singapore 728195 | 91.7 | |
| | **IGINITRFQTLL** | HLA-DPA1*02:02/DPB1*02:02 | | | |
| | IGINITRFQT-- | HLA-DPA1*02:01/DPB1*02:01 | South Africa 678570 | 83.3 | B.1.351(β) |
| | | HLA-DPA1*03:01/DPB1*23:01 | | | |
| | **SIIAYTMSLGAE** | HLA-DRB1*07:01 | | | |
| | SIIAYTMSLGVE | | South Africa 678570 | 91.7 | B.1.351(β) |
| | SIIAYTMSLGVE | | Singapore 728195 | 91.7 | |
| | **FFSNVTWFHAIH** | HLA-DPA1*02:01/DPB1*02:01 | | | |
| | FFSNVTWFHAI- | HLA-DPA1*02:02/DPB1*02:02 | Estonia 1101253 | 91.7 | B.1.1.7 (α) |
| | FFSNVTWFHAI- | HLA-DPA1*03:01/DPB1*23:01 | Italy 717978 | 91.7 | B.1.1.7 (α) |
| | FFSNVTWFHAI- | | England 728343 | 91.7 | B.1.1.7 (α) |
| | FFSNVTWFHAI- | | Botswana 6640916 | 91.7 | B.1.1.529 (o) |
| | FFSNVTWFHAI- | | Hong Kong 6530782 | 91.7 | B.1.1.529 (o) |
| | FFSNVTWFHAI- | | South Africa 6647356 | 91.7 | B.1.1.529 (o) |
| | **FKNLREFVFKNI** | HLA-DPA1*02:01/DPB1*02:01 | | | |
| | FKNLSEFVFKNI | HLA-DPA1*02:02/DPB1*02:02 | | | |
| | | HLA-DPA1*03:01/DPB1*23:01 | Brazil 804814 | 91.7 | P.1 (γ) |
| | **KIYSKHTPINLV** | HLA-DRB1*07:01 | | | |
| | KIYSKHTSINLV | | Mexico 1096985 | 91.7 | |
| | KIYSKHTPIIVR | | Botswana 6640916 | 76.9 | B.1.1.529 (o) |
| | KIYSKHTPIIVR | | Hong Kong 6530782 | 76.9 | B.1.1.529 (o) |
| | KIYSKHTPIIVR | | South Africa 6647356 | 76.9 | B.1.1.529 (o) |
| | **IKDFGGFNFSQI** | HLA-DPA1*02:02/DPB1*02:02 | | | |
| | IKDFGDFNFSQI | | Mexico 1096985 | 91.7 | |
| | IKYFGGFNFSQI | | Botswana 6640916 | 91.7 | B.1.1.529 (o) |
| | IKYFGGFNFSQI | | Hong Kong 6530782 | 91.7 | B.1.1.529 (o) |
| | IKYFGGFNFSQI | | South Africa 6647356 | 91.7 | B.1.1.529 (o) |
| | **GDSSSGWTAGAA** | HLA-DQA1*05:01/DQB1*03:01 | | | |
| | GDSSSGWTAGSA | | Australia 693279 | 91.7 | |
| | **LGAENSVAYSNN** | HLA-DQA1*05:01/DQB1*03:01 | | | |
| | LGVENSVAYSNN | | South Africa 678570 | 91.7 | B.1.351(β) |
| | LGVENSVAYSNN | | Singapore 728195 | 91.7 | |
| | **PGDSSSGWTAGA** | HLA-DQA1*05:01/DQB1*03:01 | | | |
| | PGDSSSGWTAGS | | Australia 693279 | 91.7 | |
| | **ASANLAATKMSE** | HLA-DQA1*05:01/DQB1*03:01 | | | |
| | ASANLAAIKMSE | | Brazil 804814 | 91.7 | P.1 (γ) |
| | **RSVASQSIIAYT** | HLA-DQA1*05:01/DQB1*03:01 | | | |

| | | | | | |
|---|---|---|---|---|---|
| | RSVVSQSIIAYT | | South Africa 678570 | 91.7 | B.1.351(β) |
| | **DISGINASVVNI** | HLA-DQA1*05:01/DQB1*03:01 | | | |
| | DISGINASFVNI | | Brazil 804814 | 91.7 | P.1 (γ) |
| | DISGINASFVNI | | Brazil 717944 | 91.7 | |
| | **RQIAPGQTGKIA** | HLA-DQA1*05:01/DQB1*03:01 | | | |
| | RQIAPGQTGTIA | | Brazil 804814 | 91.7 | P.1 (γ) |
| | RQIAPGQTGNIA | | South Africa 678570 | 91.7 | B.1.351(β) |
| | RQLAPGQTGNIA | HLA-B*07:02 | Botswana 6640916 | 83.3 | B.1.1.529 (o) |
| | RQLAPGQTGNIA | | Hong Kong 6530782 | 83.3 | B.1.1.529 (o) |
| | RQLAPGQTGNIA | | South Africa 6647356 | 83.3 | B.1.1.529 (o) |
| | **VGYLQPRTFLLK** | HLA-DPA1*02:02/DPB1*02:02 | | | |
| | VGYLQLRTFLLK | | Australia 693279 | 91.7 | |
| | **EGFNCYFPLQSY** | HLA-DQA1*01:01/DQB1*05:01 | | | |
| | KGFNCYFPLQSY | | Brazil 804814 | 91.7 | P.1 (γ) |
| | KGFNCYFPLQSY | | South Africa 678570 | 91.7 | B.1.351(β) |
| | KGFNCYFPLQSY | | Brazil 717944 | 91.7 | |
| | AGFNCYFPLRSY | | Botswana 6640916 | 83.3 | B.1.1.529 (o) |
| | AGFNCYFPLRSY | | Hong Kong 6530782 | 83.83 | B.1.1.529 (o) |
| | AGFNCYFPLRSY | | South Africa 6647356 | 83.3 | B.1.1.529 (o) |
| | **FSNVTWFHAIHV** | HLA-DPA1*02:02/DPB1*02:02 | | | |
| | FSNVTWFHAI-- | | Estonia 1101253 | 83.3 | B.1.1.7 (α) |
| | FSNVTWFHAI-- | | Italy 717978 | 83.3 | B.1.1.7 (α) |
| | FSNVTWFHAI— | | England 728343 | 83.3 | B.1.1.7 (α) |
| | FSNVTWFHAI-- | | Botswana 6640916 | 83.3 | B.1.1.529 (o) |
| | FSNVTWFHAI-- | | Hong Kong 6530782 | 83.3 | B.1.1.529 (o) |
| | FSNVTWFHAI-- | | South Africa 6647356 | 83.3 | B.1.1.529 (o) |
| | **GYLQPRTFLLKY** | HLA-DPA1*02:02/DPB1*02:02 | | | |
| | GYLQLRTFLLKY | | Australia 693279 | 91.7 | |
| | **MSLGAENSVAYS** | HLA-DQA1*05:01/DQB1*03:01 | | | |
| | MSLGVENSVAYS | | Singapore 728195 | 91.7 | |
| | MSLGVENSVAYS | | South Africa 678570 | 91.7 | B.1.351(β) |
| | **IEDLLFNKVTLA** | HLA-DPA1*02:02/DPB1*02:02 | Mexico 1096141 | | |
| | IEDLLFDKVTLA | | | 91.7 | |
| | **TWFHAIHVSGTN** | HLA-DQA1*05:01/DQB1*03:01 | | | |
| | TWFHAI--SGTN | | Estonia 1101253 | 83.3 | B.1.1.7 (α) |
| | TWFHAI--SGTN | | Italy 717978 | 83.3 | B.1.1.7 (α) |
| | TWFHAI—SGTN | | England 728343 | 83.3 | B.1.1.7 (α) |
| | TWFHAI--SGTN | | Botswana 6640916 | 83.3 | B.1.1.529 (o) |
| | TWFHAI--SGTN | | Hong Kong 6530782 | 83.3 | B.1.1.529 (o) |
| | TWFHAI—SGTN | | South Africa 6647356 | 83.3 | B.1.1.529 (o) |
| | **AYYVGYLQPRTF** | HLA-DRB1*01:01 | | | |
| | AYYVGYLQLRTF | | Australia 693279 | 91.7 | |
| | **GKLQDVVNQNAQ** | HLA-DRB1*04:01 | | | |
| | GKLQNVVNQNAQ | | India 3533076 | 91.7 | B.1.617.2 (δ) |
| | YLYRLFRKSNLK | HLA-DRB1*07:01 | | | |
| | YRYRLFRKSNLK | | India 3533076 | 91.7 | B.1.617.2 (δ) |
| | **PRRARSVASQSI** | HLA-DRB1*07:01 | | | |
| | PRRARSVVSQSI | | South Africa 602622 | 91.7 | |
| | HRRARSVASQSI | | Estonia 717944 | 91.7 | B.1.1.7 (α) |
| | HRRARSVASQSI | | Italy 717978 | 91.7 | B.1.1.7 (α) |
| | HRRARSVASQSI | | England 728343 | 91.7 | B.1.1.7 (α) |
| | RRRARSVASQSI | | India 3533076 | 91.7 | B.1.617.2 (δ) |
| | HRRARSVASQSI | | Botswana 6640916 | 91.7 | B.1.1.529 (o) |
| | HRRARSVASQSI | | Hong Kong 6530782 | 91.7 | B.1.1.529 (o) |
| | HRRARSVASQSI | | South Africa 6647356 | 91.7 | B.1.1.529 (o) |
| | **ARSVASQSIIAY** | HLA-DRB1*07:01 | | | |
| | ARSVVSQSIIAY | | South Africa 602622 | 91.7 | |
| | **GYFKIYSKHTPI** | HLA-DRB1*07:01 | | | |
| | GYFKIYSKHTSI | HLA-DRB1*01:01 | Mexico 1096141 | 91.7 | |
| | **NVTWFHAIHVSG** | HLA-DRB1*07:01 | | | |
| | NVTWFHAI--SG | | Estonia 1101253 | 83.3 | B.1.1.7 (α) |

| | | | | | |
|---|---|---|---|---|---|
| | NVTWFHAI--SG | | Italy 717978 | 83.3 | B.1.1.7 (α) |
| | NVTWFHAI—SG | | England 728343 | 83.3 | B.1.1.7 (α) |
| | NVTWFHAI--SG | | Botswana 6640916 | 83.3 | B.1.1.529 (ο) |
| | NVTWFHAI--SG | | Hong Kong 6530782 | 83.3 | B.1.1.529 (ο) |
| | NVTWFHAI--SG | | South Africa 6647356 | 83.3 | B.1.1.529 (ο) |
| | **QIITTDNTFVSG** | HLA-DRB1*04:01 | | | |
| | QIITTHNTFVSG | | Estonia 1101253 | 91.7 | B.1.1.7 (α) |
| | QIITTHNTFVSG | | Italy 717978 | 91.7 | B.1.1.7 (α) |
| | QIITTHNTFVSG | | England 728343 | 91.7 | B.1.1.7 (α) |
| | **NSPRRARSVASQ** | HLA-DQA1*05:01/DQB1*03:01 | | | |
| | NSHRRARSVASQ | | Estonia 1101253 | 91.7 | B.1.1.7 (α) |
| | NSHRRARSVASQ | | Italy 717978 | 91.7 | B.1.1.7 (α) |
| | NSHRRARSVASQ | | England 728343 | 91.7 | B.1.1.7 (α) |
| | NSPRRARSVVSQ | | South Africa 602622 | 91.7 | |
| | NSRRRARSVASQ | | India 3533076 | 91.7 | B.1.617.2 (δ) |
| | KSHRRARSVASQ | | Botswana 6640916 | 83.8 | B.1.1.529 (ο) |
| | KSHRRARSVASQ | | Hong Kong 6530782 | 83.8 | B.1.1.529 (ο) |
| | KSHRRARSVASQ | | South Africa 6647356 | 83.8 | B.1.1.529 (ο) |
| M | **LQFAYANRNRFL** | HLA-DRB1*07:01 | | | |
| | LQFAYANRNSFL | | Australia 710513 | 91.7 | |
| | AQFAPSASAFF- | | Brazil 804814 | 33.3 | P.1 (γ) |
| | **NRFLYIIKLIFL** | HLA-DRB1*01:01 | | | |
| | NSFLYIIKLIFL | | Australia 710513 | 91.7 | |
| | **LNTDHSSSSDNI** | HLA-DRB1*04:01 | | | |
| | LNIDHSSSSDNI | | Australia 710513 | 91.7 | |
| | **IATAMACLVGLM** | HLA-DQA1*05:01/DQB1*03:01 | | | |
| | IAIAMACLVGLM | | Australia 693279 | 91.7 | |
| | IAIAMACLVGLM | | South Africa 678570 | 91.7 | B.1.351(β) |
| | IAIAMACLVGLM | | Mexico 1096983 | 91.7 | |
| | IAIAMACLVGLM | | Malaysia 718280 | 91.7 | |
| | IAIAMACLVGLM | | Australia 710513 | 91.7 | |
| | IAIAMACLVGLM | | Estonia 717944 | 91.7 | B.1.1.7 (α) |
| | IAIAMACLVGLM | | Italy 717978 | 91.7 | B.1.1.7 (α) |
| | IAIAMACLVGLM | | England 728343 | 91.7 | B.1.1.7 (α) |
| | IAIAMACLVGLM | | Mexico 1096141 | 91.7 | |
| | IAIAMACLVGLM | | Brazil 717944 | 91.7 | |
| | IAIAMACLVGLM | | South Africa 602622 | 91.7 | |
| | IAIAMACLVGLM | | Mexico 1096985 | 91.7 | |
| | IAIAMACLVGLM | | Belgium 737393 | 91.7 | |
| | IAIAMACLVGLM | | Ecuador 660070 | 91.7 | |
| | IAIAMACLVGLM | | Saudi Arabia 490010 | 91.7 | |
| | IAIAMACLVGLM | | Germany 450204 | 91.7 | |
| | IAIAMACLVGLM | | Italy 412974 | 91.7 | |
| | IAIAMACLVGLM | | Taiwan 406031 | 91.7 | |
| | IAIAMACLVGLM | | Vietnam 408668 | 91.7 | |
| | IAIAMACLVGLM | | Wuhan 402125 | 91.7 | |
| | IAIAMACLVGLM | | Wuhan 406798 | 91.7 | |
| | IAIAMACLVGLM | | Ecuador 728204 | 91.7 | |
| | IAIAMACLVGLM | | New Zealand 456378 | 91.7 | |
| | IAIAMACLVGLM | | Singapore 728195 | 91.7 | |
| | IAIAMACLVGLM | | USA 6142566 | 91.7 | |
| | SRIGMEVTPSGTM | | Brazil 804814 | 5.3 | P.1 (γ) |
| | **ANRNRFLYIIKL** | HLA-DPA1*02:02/DPB1*02:02 | | | |
| | ANRNSFLYIIKL | | Australia 710513 | 91.7 | |
|  | SASAFF------ |  | Brazil 804814 | 8.3 | P.1 (γ) |
| | **NYKLNTDHSSSS** | HLA-DRB1*04:01 | | | |
| | NYKLNIDHSSSS | | Australia 693279 | 91.7 | |
| N | **IAQFAPSASAFF** | HLA-DRB1*07:01 | | | |
| | IALFAPSASAFF | | Mexico 1096985 | 91.7 | |
| | **GIIWVATEGALN** | HLA-DRB1*01:01 | | | |
| | GIIWVATEGAFN | | South Africa 602622 | 91.7 | |
| | **YFYYLGTGPEAGL** | HLA-DRB1*01:01 | | | |

| | | | | | |
|---|---|---|---|---|---|
| | YFYYLGTGPE**S**GL | | Brazil 717944 | 91.7 | |
| | **AIVLQLPQGTTL**<br>AIVLQLPQGT**I**L | HLA-DRB1*01:01 | <br>Ecuador 660070 | <br>91.7 | |
| | **LALLLLDRLNQL**<br>L**V**LLLLDRLNQL<br>L**V**LLLLDRLNQL | HLA-DRB1*04:01 | <br>Belgium 737393<br>Australia 693279 | <br>91.7<br>91.7 | |
| | **IIWVATEGALNT**<br>IIWVATEGA**F**NT | HLA-DRB1*01:01 | <br>South Africa 602622 | <br>91.7 | |
| | **LDRLNQLESKMS**<br>LDRLNQLESKM**F**<br>LDRLNQLESKM**F**<br>LDRLNQLESKM**F**<br>LDRLNQLESK**I**S<br>LDRLNQLESK**I**S | HLA-DRB1*04:01 | <br>Estonia 717944<br>Italy 717978<br>England 728343<br>Brazil 717944<br>Ecuador 728204 | <br>91.7<br>91.7<br>91.7<br>91.7<br>91.7 | <br>B.1.1.7 (α)<br>B.1.1.7 (α)<br>B.1.1.7 (α) |
| | **DAALALLLLDRL**<br>DAAL**V**LLLLDRL<br>DAAL**V**LLLLDRL | HLA-DPA1*02:02/DPB1*02:02 | <br>Belgium 737393<br>Australia 693279 | <br>91.7<br>91.7 | |
| | **TGPEAGLPYGAN**<br>TGPE**S**GLPYGAN | HLA-DQA1*05:01/DQB1*03:01 | <br>Brazil 717944 | <br>91.7 | |
| | **RITFGGPSDSTG**<br>RITFGGPSDS**I**G | HLA-DQA1*05:01/DQB1*03:01 | <br>Mexico 1096985 | <br>91.7 | |
| | **QIAQFAPSASAF**<br>QIA**L**FAPSASAF | HLA-DQA1*05:01/DQB1*03:01 | Mexico 1096985 | <br>91.7 | |
| | **AGNGGDAALALL**<br>AGNGGDAAL**V**LL<br>AGNGGDAAL**V**LL<br>AGNG**C**DAALALL | HLA-DQA1*05:01/DQB1*03:01 | <br>Belgium 737393<br>Australia 693279<br>India 3533076 | <br>91.7<br>91.7<br>91.7 | <br><br><br>B.1.617.2 (δ) |
| | **IWVATEGALNTP**<br>IWVATEGA**F**NTP | HLA-DRB1*01:01 | <br>South Africa 602622 | <br>91.7 | |
| | **RMAGNGGDAALA**<br>RMAGNGGDAAL**V**<br>RMAGNGGDAAL**V**<br>**I**MAGNGGDAALA<br>**I**MAGNGGDAALA<br>R**I**AGNGGDAALA<br>RMAGNG**C**DAALA | HLA-DQA1*05:01/DQB1*03:01 | <br>Australia 710513<br>Belgium 737393<br>Mexico 1096142<br>Malaysia 718280<br>Ecuador 660070<br>India 3533076 | <br>91.7<br>91.7<br>83.3<br>83.3<br>83.3<br>91.7 | <br><br><br><br><br><br>B.1.617.2 (δ) |
| | **SSRGTSPARMAG**<br>SSRGT**F**PARMAG<br>SSRG**I**SPARMAG<br>SSRG**I**SPA**I**MAG<br>SS**KR**TSPA**I**MAG<br>SS**KR**TSPARMAG<br>SS**KR**TSPARMAG<br>SS**KR**TSPARMAG<br>SS**KR**TSPARMAG<br>SS**KR**TSPARMAG<br>SS**KR**TSPARMAG<br>SS**KR**TSPARMAG<br>SS**KR**TSPARMAG<br>SS**KR**TSPARMAG<br>SS**M**GTSPARMAG | HLA-DQA1*05:01/DQB1*03:01 | Australia 710513<br>Belgium 737393<br>Ecuador 660070<br>South Africa 678570<br>Mexico 1096142<br>Malaysia 718280<br>Australia 693279<br>Estonia 717944<br>Italy 717978<br>England 728343<br>Mexico 1096141<br>Brazil 717944<br>Brazil 804814<br>South Africa 602622<br>India 3533076 | <br>83.3<br>83.3<br>83.3<br>75<br>83.3<br>83.3<br>83.3<br>83.3<br>83.3<br>83.3<br>83.3<br>83.3<br>83.3<br>91.7 | <br><br><br>B.1.351(β)<br><br><br><br>B.1.1.7 (α)<br>B.1.1.7 (α)<br>B.1.1.7 (α)<br><br><br>P.1 (γ)<br><br>B.1.617.2 (δ) |

**-In red the amino acid changes between the Wuhan sequence and the sequences of other countries.**

**Table 8. SARS-CoV-2 reported linear B cell epitopes identity (%) shared with other coronaviruses**

| Protein | SARS-CoV-2 | SARS | MERS | 229E | HKU1 | NL63 | OC43 |
|---|---|---|---|---|---|---|---|
| S | DPFLGVYYHKNNKSWME | 17.6 | 1.9 | 2.4 | 15.6 | 8.1 | 13.8 |
| | MDLEGKQGNFKNL | 53.8 | 0 | 13.3 | 0 | 13.3 | 0 |
| | KHTPINLVRDLPQGFS | 56.2 | 6.2 | 5.9 | 5.6 | 4.5 | 12.5 |
| | GDEVRQIAPGQTGKIADYNYKLPDD | 92 | 20.7 | 2.4 | 20 | 5.9 | 23.1 |
| | SNKKFLPF | 62.5 | 0 | 50 | 7.7 | 25 | 7.7 |
| | NCTEVPVAIHADQLTPT | 64.7 | 3.6 | 6.2 | 23.5 | 3.1 | 29.4 |
| | RVYSTGSNVFQ | 81.8 | 0 | 9.1 | 9.1 | 9.1 | 18.2 |
| | VNNSYECDIPI | 81.8 | 21.4 | 6.7 | 12.5 | 6.7 | 15.4 |
| | ASYQTQTNSPRRARSVASQ | 31.6 | 40 | 6.5 | 24 | 15.8 | 15.8 |
| | YTMSLGAENSVAYSNN | 81.2 | 11.1 | 5 | 8 | 5 | 3.4 |
| | KQIYKTPPIKDFGGF | 80 | 40 | 20 | 3.2 | 13.3 | 15 |
| | LADAGFIKQYGDCLG | 86.7 | 35.3 | 20 | 46.7 | 18.8 | 46.7 |
| | RNFYEPQIITTD | 83.3 | 41.7 | 41.7 | 25 | 16.7 | 33.3 |
| | SGTNGTKRFDN | 27.3 | 26.7 | 0 | 12.5 | 4.3 | 6.2 |
| | NGTITD | 100 | 33.3 | 9.1 | 50 | 9.1 | 33.3 |
| | YQAGSTPCNGV | 20 | 21.4 | 0 | 13.3 | 0 | 25 |
| | YGFQPTNGVGYQ | 66.7 | 5 | 8.3 | 10.5 | 6.7 | 5.3 |
| | TVCGPKKSTN | 80 | 20 | 8.3 | 17.6 | 20 | 23.1 |
| | RDIADTTDAVRDPQ | 64.3 | 4 | 4 | 4.2 | 0 | 8.3 |
| | VEQDKNTQE | 66.7 | 22.2 | 6.7 | 0 | 8.3 | 5.9 |
| | ILPDPSKPSKRS | 83.3 | 20 | 22.2 | 26.7 | 25 | 31.2 |
| | DSLSST | 50 | 16.7 | 16.7 | 33.3 | 50 | 16.7 |
| | PAQEKNFTT | 77.8 | 11.1 | 33.3 | 16.7 | 33.3 | 8.3 |
| | KNHTSPDVDLG | 100 | 27.3 | 28.6 | 38.5 | 7.1 | 54.5 |
| | **GQSKRVDFC** | 100 | 66.7 | 66.7 | 55.6 | 55.6 | 55.6 |
| M | LNTDHSSSSD | 70 | 7.7 | 13.3 | 30 | 5.6 | 18.2 |
| | PLLESE | 73.3 | 50 | 16.7 | 33.3 | 16.7 | 33.3 |
| | YRIGNYKLNTDHSSSSDNIA | 85 | 21.7 | 14.3 | 35 | 10 | 30 |
| N | FGGPSDSTGSNQNGERSGARSKQRRPQGLPNN | 81.2 | 31.2 | 15.6 | 24.3 | 2.4 | 19.5 |
| | SKQLQQSMSSADS | 100 | 0 | 5.6 | 4.2 | 5.6 | 5.9 |
| | RIRGGDGKMKDL | 83.3 | 41.7 | 41.7 | 30.8 | 38.5 | 23.1 |
| | TGPEAGLPYGANK | 92.3 | 69.2 | 23.1 | 46.2 | 30.8 | 46.2 |
| | SKMSGKGQQQQGQTVTKKSAAEASKKPRQKRTATKAYN | 94.7 | 50 | 18.6 | 38.1 | 25 | 35.7 |
| | TDYKHW | 100 | 33.3 | 11.1 | 16.7 | 33.3 | 14.3 |
| | KLDDKDPNFKD | 90 | 60 | 10 | 30 | 23.1 | 20 |

**-Epitope greater than 50% shared with other coronaviruses is shown in bold**

**Table 9. SARS-CoV-2 reported T cell epitopes identity (%) shared with other coronaviruses**

| | Protein | SARS-CoV-2 | SARS | MERS | 229E | HKU1 | NL63 | OC43 |
|---|---|---|---|---|---|---|---|---|
| MHC-I | S | QPTESIVRF | 44.4 | 33.3 | 30 | 22.2 | 9.1 | 33.3 |
| | | YQDVNCTEV | 88.9 | 22.2 | 7.1 | 33.3 | 0 | 22.2 |
| | | **IPTNFTISV** | 77.8 | 66.7 | 66.7 | 77.8 | 66.7 | 55.6 |
| | | RFDNPVLPF | 77.8 | 15.4 | 0 | 16.7 | 25 | 25 |
| | | VFKNIDGYF | 66.7 | 5.9 | 33.3 | 6.2 | 40 | 6.2 |
| | | FPREGVFVS | 88.9 | 33.3 | 6.2 | 33.3 | 6.2 | 44.4 |
| | | LEPLVDLPI | 44.4 | 22.2 | 18.2 | 5.9 | 18.2 | 5.9 |
| | | TPCNGVEGFNCY | 41.7 | 17.6 | 0 | 3.2 | 8.3 | 4.5 |
| | | YQPYRVVVL | 100 | 6.7 | 6.2 | 3.8 | 16.7 | 4.8 |
| | | NATRFASVYAWNRK | 78.6 | 9.1 | 18.8 | 35.7 | 18.8 | 35.7 |
| | M | SSDNIALLV | 88.9 | 33.3 | 30 | 44.4 | 30 | 44.4 |
| | N | KHWPQIAQF | 100 | 55.6 | 44.4 | 22.2 | 44.4 | 22.2 |
| | | LPNNTASWF | 100 | 66.7 | 8.3 | 27.3 | 6.7 | 33.3 |
| | | AEGSRGGSQASSRSSSR | 100 | 56.8 | 21.1 | 41.2 | 38.1 | 47.1 |
| MHC-II | S | MFVFLVLLPLVSSQCVNLT | 34.8 | 20.6 | 0 | 19.2 | 21.1 | 13.9 |
| | | LDSKTQSLLIVN | 41.7 | 3.3 | 3.4 | 4.5 | 4.1 | 7.1 |
| | | RQIAPGQTGKIA | 91.7 | 16.7 | 5 | 16.7 | 5.3 | 13.3 |
| | | LQSYGFQPTNGVG | 53.8 | 23.1 | 5 | 15 | 5.3 | 10 |
| | | KKSTNLVKNKCV | 66.7 | 28.6 | 33.3 | 25 | 41.7 | 33.3 |
| | | GLTGTGVLTESN | 83.3 | 33.3 | 41.7 | 41.7 | 41.7 | 41.7 |
| | | TPCSFGGVSVITPGTN | 93.8 | 31.2 | 25 | 31.2 | 29.4 | 25 |
| | | TWRVYSTGSNVFQTRAGCLIGAE | 82.6 | 20 | 2.9 | 17.4 | 11.5 | 21.7 |
| | | VKQIYKTPPIKDFGGFNFSQILPDPSKSK | 83.3 | 31.4 | 23.5 | 15 | 20.6 | 25 |
| | M | **LRGHLRIAGHHL** | 75 | 50 | 50 | 58.3 | 41.7 | 58.3 |
| | N | PRITFGGPSDSTGSN | 80 | 26.7 | 4 | 6.7 | 4.5 | 13.3 |
| MHC-I & MHC-II | S | GVYFASTEKSNI | 75 | 10.3 | 0 | 9.5 | 2.1 | 14.3 |
| | | NVVIKVCEFQFCNDPFL | 52.9 | 3.6 | 20 | 21.1 | 0 | 23.5 |
| | | SANNCTFEYVSQPFLMDLEGKQGN | 54.2 | 22.2 | 2.6 | 20 | 5 | 16 |
| | | CVADYSVLYNSASFSTFKCYGVSPTKLN | 92.9 | 25 | 14.3 | 21.4 | 14.3 | 21.4 |
| | | DLCFTNVYADSFVI | 85.7 | 21.4 | 4.5 | 35.7 | 5.6 | 35.7 |
| | | NYNYLYRLFRKSNLKPFERDISTEIYQ | 52 | 19.4 | 3.8 | 8.2 | 9.5 | 7.8 |
| | | SFIEDLLFNKVTLADAGFIK | 95 | 65 | 40 | 65 | 45 | 65 |
| | | GWTFGAGAALQIPFAMQMAYRFNGIGVTQNVLYENQK | 100 | 56.8 | 35.1 | 48.6 | 35.1 | 48.6 |
| | | QKLIANQFNSAIGKIQDSLSSTAS | 62.5 | 50 | 21.1 | 54.2 | 28.9 | 54.2 |
| | | ALGKLQDVVNQNAQALNTLVKQLSSNFGAISSVLNDILSRL | 100 | 46.3 | 48.8 | 56.1 | 48.8 | 53.7 |
| | | RLDKVEAEVQIDRLITGRLQSLQTYVTQQLIRAAEIRASANLAATKM | 100 | 51.1 | 48.9 | 53.2 | 48.9 | 53.2 |
| | | QSAPHGVVFLHVTVPAQEK | 85 | 40 | 50 | 45 | 45 | 45 |
| | | FVSNGTHWFVTQRNFYEPQII | 71.4 | 34.6 | 26.1 | 33.3 | 25 | 33.3 |
| | | LDKYFKNHTSPDVDLGDISGINASVVNIQKEIIDRLNEVAKNL | 97.6 | 31 | 18.6 | 34.1 | 19 | 43.2 |
| | M | GTITVEELK | 88.9 | 7.1 | 15.4 | 6.7 | 11.1 | 6.7 |
| | | LKKLLEQWNLVIGFLFLTWICLLQFAYANRNRFLYIIKLIFLWLLWPVTLACF | 90.6 | 32.1 | 34 | 39.6 | 31.6 | 41.5 |
| | | FLWLLWPVTLACFVLAAVYRINWITGGIAIAMACLVGLMWLSYFIASFRLFARTRSMWSFNP | 95.7 | 37.7 | 42.3 | 43.5 | 39.2 | 44.9 |
| | | RCIKDLPKEITVATSRTLSYYKLGASQRVAG | 93.8 | 34.4 | 29.4 | 27.3 | 32.4 | 33.3 |
| | | DSGFAAYSRY | 90 | 60 | 30 | 40 | 40 | 50 |
| | | RYRIGNYKL | 100 | 55.6 | 22.2 | 44.4 | 22.2 | 44.4 |
| | N | PDDQIGYYRRATRRIRGGDGKM | 95.5 | 40.9 | 31.8 | 30.4 | 39.1 | 30.4 |
| | | GIIWVATEGALNTPKDHI | 94.4 | 38.9 | 38.9 | 42.1 | 45 | 42.1 |
| | | GTRNPANNAAIVLQLPQGTTLPKGFYA | 92.6 | 59.3 | 23.3 | 37 | 14.7 | 40.7 |
| | | RMAGNGGDAALALLLLDRLNQLESKMSG | 82.1 | 34.3 | 21.6 | 15.8 | 25.7 | 21.2 |
| | | QQQQGQTVTKKSAAEASKK | 100 | 44.4 | 5.9 | 27.3 | 10.7 | 27.3 |
| | | WPQIAQFAPSASAFFGMSRIGMEVTPSGTWL | 100 | 33.3 | 30.3 | 29.3 | 27.3 | 31.8 |
| | | NFKDQVILLNKHIDAYKTFPPTEPKKD | 92.6 | 48.1 | 3.3 | 15.6 | 21.4 | 21.4 |

**-Epitopes greater than 50% shared with other coronaviruses are shown in bold**

**Bibliography (Tables)**

1. Denis M, Vandeweerd V, Verbeeke R, Laudisoit A, Reid T, Hobbs E, et al. Covipendium: Information available to support the development of medical countermeasures and interventions against COVID-19. Transdisciplinary Insights. 2021 Mar 12;4(1):1–296.
2. Kumar S. Drug and vaccine design against Novel Coronavirus (2019-nCoV) spike protein through Computational approach. Preprints [Internet]. 2020; Available from: www.preprints.org
3. Singh Slathia P, Sharma P. Prediction of T and B cell epitopes in the proteome of SARS-CoV-2 for potential use in diagnostics and vaccine design. ChemRxiv [Internet]. 2020; Available from: https://www.who.int/emergencies/diseases/novel-coronavirus-2019
4. Polyiam K, Phoolcharoen W, Butkhot N, Srisaowakarn C, Thitithanyanont A, Auewarakul P, et al. Immunodominant linear B cell epitopes in the spike and membrane proteins of SARS-CoV-2 identified by immunoinformatics prediction and immunoassay. Scientific Reports 2021 11:1 [Internet]. 2021 Oct 14 [cited 2021 Nov 23];11(1):1–17. Available from: https://www.nature.com/articles/s41598-021-99642-w
5. le Bert N, Tan AT, Kunasegaran K, Tham CYL, Hafezi M, Chia A, et al. SARS-CoV-2-specific T cell immunity in cases of COVID-19 and SARS, and uninfected controls. Nature. 2020 Aug 20;584(7821):457–62.
6. Ahmed SF, Quadeer AA, McKay MR. COVIDep: a web-based platform for real-time reporting of vaccine target recommendations for SARS-CoV-2. Nature Protocols. 2020 Jul 1;15(7):2141–2.
7. Amrun SN, Lee CYP, Lee B, Fong SW, Young BE, Chee RSL, et al. Linear B-cell epitopes in the spike and nucleocapsid proteins as markers of SARS-CoV-2 exposure and disease severity. EBioMedicine [Internet]. 2020 Aug 1 [cited 2021 Nov 23];58:102911. Available from: http://www.thelancet.com/article/S2352396420302863/fulltext
8. Grifoni A, Sidney J, Zhang Y, Scheuermann RH, Peters B, Sette A. A Sequence Homology and Bioinformatic Approach Can Predict Candidate Targets for Immune Responses to SARS-CoV-2. Cell Host and Microbe. 2020 Apr 8;27(4):671-680.e2.
9. Qamar MT ul, Shahid F, Ashfaq UA, Aslam S, Fatima I, Fareed MM, et al. Structural modeling and conserved epitopes prediction against SARS-COV-2 structural proteins for vaccine development. Research Squeare. 2020;
10. Peng Y, Mentzer AJ, Liu G, Yao X, Yin Z, Dong D, et al. Broad and strong memory CD4+ and CD8+ T cells induced by SARS-CoV-2 in UK convalescent individuals following COVID-19. Nature Immunology. 2020 Nov 1;21(11):1336–45.
11. Shomuradova AS, Vagida MS, Sheetikov SA, Zornikova K v., Kiryukhin D, Titov A, et al. SARS-CoV-2 Epitopes Are Recognized by a Public and Diverse Repertoire of Human T Cell Receptors. Immunity. 2020 Dec 15;53(6):1245-1257.e5.

20. Fast E, Altman RB, Chen B. Potential T-cell and B-cell epitopes of 2019-nCoV. bioRxiv. 2020;

21. Kar T, Narsaria U, Basak S, Deb D, Castiglione F, Mueller DM, et al. A candidate multi-epitope vaccine against SARS-CoV-2. Scientific Reports. 2020 Dec 1;10(1).

22. Schreibing F, Hannani M, Ticconi F, Fewings E, Nagai JS, Begemann M, et al. Dissecting CD8+ T cell pathology of severe SARS-CoV-2 infection by single-cell epitope mapping. BioRxiv [Internet]. 2021 [cited 2021 Nov 23]; Available from: https://biorxiv.org/cgi/content/short/2021.03.03.432690
